# Evolutionary and ecological determinants of the phenology of births in wild large herbivores, a systematic review

**DOI:** 10.64898/2025.12.14.694258

**Authors:** Lucie Thel, Christophe Bonenfant, Simon Chamaillé-Jammes

## Abstract

**Introduction:** Timing of birth is a major determinant of individuals’ reproductive success and phenotypic performances, driven by multiple factors. However, there is currently no quantitative assessment of the support received by the drivers of the phenology of births in mammals.

**Aims:** our objective was to identify the drivers tested for their effect on the phenology of births in large herbivores and how much empirical support they received, as well as determine thematic, taxonomic and spatial gaps in the literature.

**Methods:** We conducted a systematic review focusing on the drivers of the phenology of births in large herbivores. We extracted the outcomes of 124 articles and assessed the reliability of their findings using a general index based on the quality of the data and methods used.

**Results and Discussion:** We identified a major research gap in Asian and South and Central American species. The effect of population and individual characteristics remained marginally accounted for, even though there was evidence of their importance in the determination of the phenology of births. The two dominant hypotheses (seasonality and predation) received good to moderate empirical support regarding their effect on birth timing and synchrony, respectively. Given the strong geographic and methodological biases, these hypotheses should be considered as most consistently supported in the existing literature rather than definitively established across systems.

**Conclusions:** We encourage research in these understudied domains, as well as large scale multi-species comparative analyses to fill in these gaps and improve our understanding of reproductive phenology.

**Graphical Abstract:** 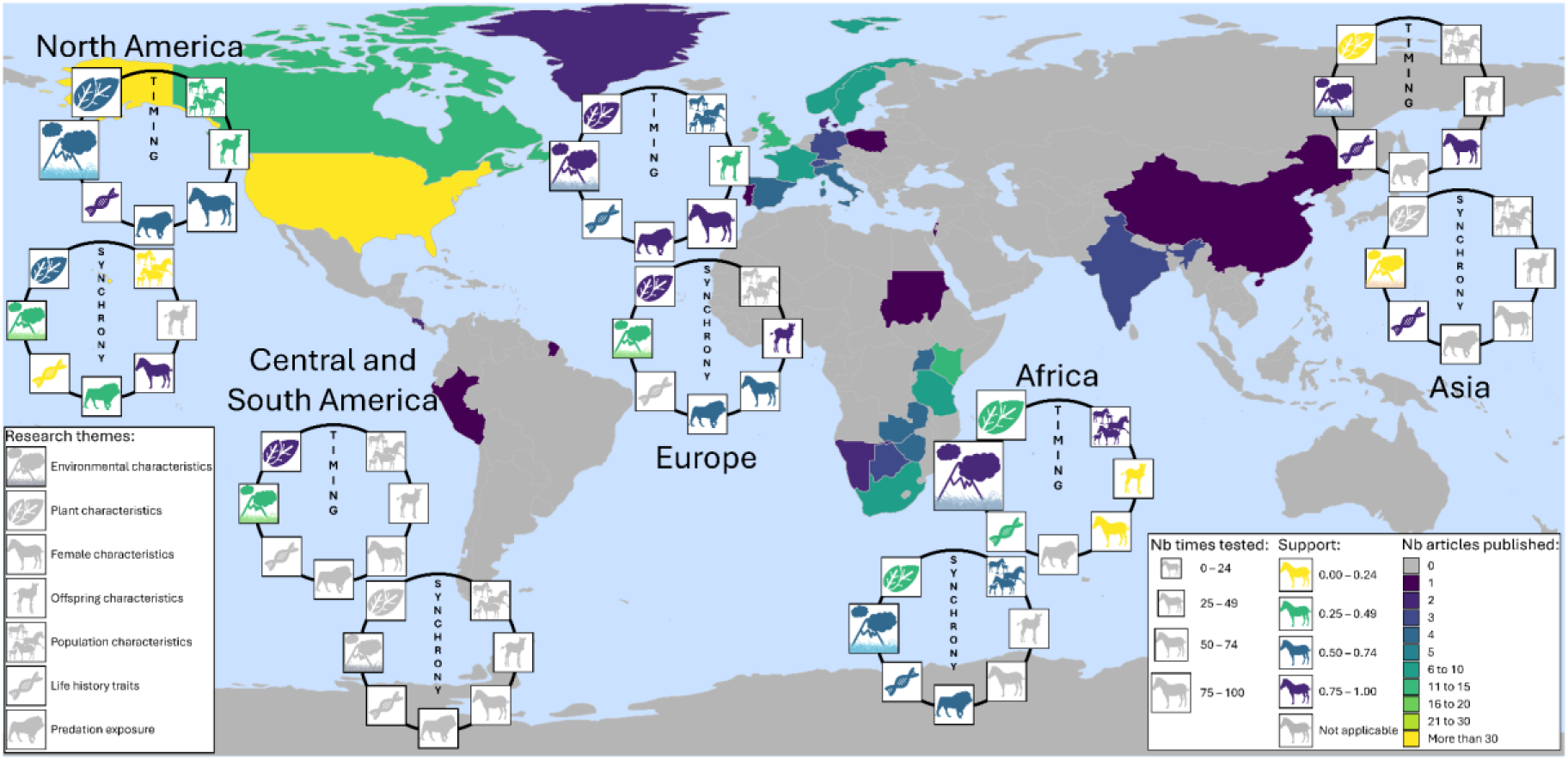

This semi-systematic review supports the two dominant drivers of birth phenology, the seasonality and predation hypotheses. Even though there is evidence of their importance, the effects of female, offspring and population characteristics remain marginally accounted for. Asian and South and Central American species are currently understudied.

## Introduction

The timing of birth is a major determinant of an individual’s reproductive success, as it may affect both the short- and long-term phenotypic performance of the offspring (e.g., via early growth, Côté and Festa-Bianchet 2001; via density-dependent phenomena, Williams et al. 2013), the survival of the parents, and possibly the later reproductive success of both the parents and the offspring (Green and Rothstein 1993a). The timing of birth can be affected by numerous drivers, such as the seasonality of the environment (e.g., food availability, Bunnell 1982; temperature variations, McNutt et al. 2019; date of snowmelt, Laforge et al. 2023), interacting with life history traits (e.g., diet, Sinclair et al. 2000; mating system, Boness et al. 1995; sociality, Bertram 1975) and predation pressure (O’Donoghue and Boutin 1995). For instance, most mammal species in the northern hemisphere give birth in spring when the vegetation blooms, providing sufficient food of high quality to cope with the energetic costs of gestation and lactation (e.g., in the roe deer (*Capreolus capreolus*), Plard et al. 2014), but there is evidence that individuals can also actively tune their parturition date by adjusting their gestation length in response to variations of temperature and snow conditions during the ongoing year (e.g., in the reindeer (*Rangifer tarandus*), Paoli et al. 2020). The phenology of births in a population is thus the result of a complex interaction between multiple ecological and biological drivers acting at different spatio-temporal scales, which induce both plastic and evolutionary responses (Radchuk et al. 2019).

The study of the phenology of births consists mainly in describing the birth timing (when births happen) and synchrony (how spread out births are over time) in a given population and understanding their drivers (Thel et al. 2022). Although some main patterns of animal phenology have been known and described for a long time, timing and synchrony of births are highly variable in time and space (Bronson 1989). As a matter of fact, the phenology of births in mammals ranges from year-round breeding (e.g., the chacma baboon (*Papio ursinus*) in Tsaobis Nature Park, Namibia, Dezeure et al. 2021) to a concentration of births during a few weeks (e.g., 80% of births happen in less than 30 days in a French population of roe deer (Gaillard et al. 1993). The phenology of births varies from one species to the other in the same location (e.g., in the guild of East African antelopes in the Serengeti, Tanzania, Sinclair et al. 2000), but also from one population to the other within the same species across its geographical range. In Africa, the impala (*Aepyceros melampus*) gives birth year round in Kenya and Tanzania, but seasonally in Zambia, Botswana and Rwanda (Ansell 1960, Dasmann and Mossman 1962, Leuthold and Leuthold 1975, Sinclair et al. 2000, Moe et al. 2007).

What factors drive the phenology of births has been a major research avenue, producing numerous studies going back a long way in time (e.g., in large herbivores: Ansell 1960, Spinage 1973, Bunnell 1980, Gaillard et al. 1993, Post et al. 2003, Ryan et al. 2007, Ogutu et al. 2010, Paoli et al. 2018, Neumann et al. 2020) and generating several qualitative and quantitative reviews and book chapters (e.g., Rutberg 1987, Bronson 1989, Post 2003, Brockman and van Schaik 2005, Zerbe et al. 2012, Heldstab et al. 2018). To account for such a variability of phenological patterns, the main and most frequently advanced hypothesis is the seasonality hypothesis according to which the timing of births is primarily set by the seasonality of climate and food resources (Bunnell 1980, Festa-Bianchet 1988, Post 2003). The timing of births is expected to match with environmental conditions determining the best period of the year to give birth to cope with the costs of reproduction, thus improving parent’s reproductive success (Green and Rothstein 1993a), and to reduce climate-related mortality for the newborn such as hypothermia (e.g., in the Eurasian lynx (*Lynx lynx*), Mattisson et al. 2022). The length of the favorable season may also shape the synchrony of births, whereby shorter favorable windows, such as at high latitude or altitude, favor more synchronous births (Bunnell 1982, Heldstab et al. 2018). Theory also predicts more synchronous births where environmental cues are predictable (Mattisson et al. 2022). Nevertheless, despite being intrinsically related, timing and synchrony of births do not necessarily respond to the same selection pressures (Ahrestani et al. 2011).

In addition to its sensitivity to environmental constraints, synchrony of births can also be shaped by other drivers such as social factors and, in prey species such as large herbivores and rodents, predation pressure. The predation hypothesis states that the synchrony of births is shaped by the type of predation and the anti-predator behaviour of the juvenile (Estes 1976, Rutberg 1987, Ims 1990a, Ims 1990b, O’Donoghue and Boutin 1995). Predation risk increases for juveniles born very early or late in the season because very few other newborns are available to dilute their own predation risk. Predation can hence produce stabilizing selection toward a central birth date in some populations (Loe et al. 2005). Anti-predator strategies of the juvenile, such as the hider-follower gradient in large herbivores (Lent 1974) can modulate this dilution effects, favouring either high birth synchrony to swamp predators, or spreading of births to reduce risks of being preyed upon (Estes 1976, Sinclair et al. 2000). Predator satiation benefits seem strongest in follower species whose young follow their mother from birth, where large aggregations create a confusion or dilution effect. The theory suggests that high synchrony of births could be less beneficial with the hider anti-predator strategy, which consists in hiding the young for several days to weeks after birth. In highly social species, intra-specific interactions can also play a major role in determining the level of synchrony between the members of the group. For instance, small mammals displaying communal breeding such as the banded mongoose (*Mungos mungo*) tend to breed synchronously to limit the risk of infanticide and competition between offspring (Hodge et al. 2011). In a pride of lions (*Panthera leo*), females tend to become fertile independently of the time of the year, but rather when a new male takes over the pride and kills the offspring sired by the previous male, leading to synchronous births (Bertram 1975).

Over the last century, a growing number of more specific hypotheses have been put forward to account for the deviation of birth phenology from the general matching with environmental seasonality (e.g., Zerbe et al. 2012, Heldstab et al. 2021). For instance, it is generally accepted that grazer species should have a shorter birth period than browser species because the availability of their food resource, mainly composed of grasses, is more concentrated in time than tree-based resources such as leaves, preferred by browsers (diet hypothesis, Leuthold and Leuthold 1975). However, empirical tests conducted on some populations led to inconclusive outcomes or results contradicting theoretical expectations (e.g., Rutberg (1987), English et al. (2012) and Ogutu et al. (2014) found no support for the diet hypothesis; Michel et al. (2020) found that synchronising birth also seems beneficial to a hider species). More generally, current reviews on the phenology of births are either dated and do not include the latest publications on the subject (e.g., Rutberg 1987), they focus on a limited geographical range (e.g., Bunnell 1982), or they are based mainly on captive individuals (e.g., Zerbe et al. 2012). Currently, there is no quantitative assessment of the reliability and empirical validity of the hypotheses tested in terms of phenology of births in mammals. A comprehensive summary of the methodological approaches and robustness of the tests used to explore these different hypotheses is also currently lacking.

Due to their great diversity in traits and environments they live in, as well as their pivotal position in the ecological processes as primary consumers and prey species, large herbivores (see definition in the Methods section) provide an ideal framework for analysing and understanding the regulatory processes of the phenology of births in mammals. Large herbivores are present at almost all latitudes from the poles (e.g., the caribou (*Rangifer tarandus*), Post et al. 2003) to the equator (e.g., the Bornean yellow muntjac (*Muntiacus atherodes*), Giman et al. 2007), and occur in habitats ranging from sea level (e.g., the moose (*Alces alces*), Singh et al. 2012) to mountainous landscapes as elevated as 5000 m a.s.l. (e.g., the alpine musk deer (*Moschus chrysogaster*), Buzzard et al. 2018). Those species present a large variety of life history traits in terms of their diet (Hofmann 1989), sociality (Szemán et al. 2021), and reproductive traits (e.g., gestation length, Owen-Smith and Ogutu 2013; mating system, Bowyer et al. 2020), amongst others. Additionally, they are exposed to a wide range of predation pressures, from the complete absence of predators, as it has been the case for almost two centuries in Rum Island, Scotland (Shi et al. 2010), to the full guild of medium and large predators such as in the Serengeti ecosystem, Tanzania (Sinclair and Northon-Griffiths 1995), including recovering systems such as Yellowstone National Park, United States of America (Ripple et al. 2010). Such a diversity of conditions and traits gathered in one clade has generated a vast range of research and participated in the emergence and development of major ecological theories revolving around reproductive biology (e.g., density-dependence effects, Bonenfant et al. 2009; capital-income breeding strategies, Apollonio et al. 2020), some of them directly related to the phenology of births (e.g., Gosling 1969, Post 2003).

In this study, we conducted a semi-systematic review of the scientific literature focusing on the drivers of the phenology of births in large herbivores living in natural conditions. We identified 124 relevant scientific articles published since the early 1960’s, from which we summarised the different hypotheses and how much empirical support they have received so far. We highlighted the hypotheses, the geographic areas and the taxonomic groups that are currently understudied and we suggested new avenues of research for a better understanding of the drivers of the phenology of births in large herbivores, and in mammals in general.

## Methods

### Focus of the literature search

The “ungulates” consist of a polyphyletic group generally used to describe herbivorous hoofed mammals with a body mass > 2 kg. It commonly includes the orders Artiodactyla (i.e., even number of fingers) and Perissodactyla (i.e., uneven number of fingers), but can be extended to some large marsupials and to the Proboscidea (i.e., characterised by a trunk) according to the definition selected (Danell et al. 2006). As part of the Artiodactyla, the Suina (i.e., pigs and peccaries) and the Cetacea (i.e., aquatic mammals), which are not strictly herbivorous, can also be included in the ungulates *lato sensu*. Based on their similarities in terms of eco-evolutionary traits, we retained in this semi-systematic review the Artiodactyla (including the Suina but not the Cetacea), the Perissodactyla and the Proboscidea, and refer to them as “large herbivores”.

### Systematic literature search

We conducted an initial literature search (16^th^ of July, 2025) on the Web of Science (www.webofscience.com, WoS hereafter) using all databases available (Web of Science Core Collection, Grants Index, Inspec, KCI-Korean Journal Database, Medline, Preprint Citation Index, ProQuest Dissertations & Theses Citation Index, SciELO Citation Index). We searched all scientific articles, review articles, books and data papers written in English and published before 2025. We excluded the year 2025 from our review to avoid partial representation of the year in terms of number of publications. We included in the following catalogues (inbuilt in the WoS search engine): “reproductive biology”, “environmental science ecology”, “zoology”, “behavioural sciences”, “biodiversity conservation”, “demography”, “evolutionary biology” and “remote sensing”. To conduct the search, we used the following list of keywords: “phenolog* OR *synchron* OR timing* OR period* OR *season* OR date* OR pattern*” (“OR” allowed us to search for records containing any of the terms separated by the operator; “*” allowed us to search for words with variant zero to many characters, e.g., the use of “period*” allowed us to search for words such as “period”, “periods”, “periodic”, “periodicity”). We combined these terms with the Boolean connector “AND” (“AND” finds records containing both terms separated by the operator) with the following list of keywords: “breed* OR birth* OR parturition* OR reproduc* OR calv* OR fawn* OR lamb* OR foal* OR farrow* OR litter* OR offspring* OR newborn*”, and looked for this combination of keywords in the title of the articles referenced in the WoS.

We obtained n = 15919 initial results (see PRISMA diagram in Supplementary Material 1, Page et al. 2021). We excluded n = 28 duplicates based on identical DOI and/or identical combination of title and list of authors. Based on a quick screening of the title (and abstract, when necessary), we excluded n = 15822 irrelevant articles that were either not dealing with large herbivores reproduction, or when dealing with large herbivores reproduction, that were focusing on: i) populations subject to experimental designs or veterinary treatments; ii) populations in captive, farming or zoo conditions; iii) exotic or feral populations; iv) subjects not dealing with phenology of births in general (e.g., male reproduction, mating instead of parturition phenology). We also excluded purely methodological articles, which presented methods to estimate the date of birth but did not explore the effect of any driver on the phenology of births. We extracted n = 69 articles dealing explicitly with the phenology of births in large herbivores in wild conditions. Although the WoS references publications since 1637, we mostly had access to publications after 1993 for some databases due to academic subscription restrictions, and we acknowledge that due to probable imperfect referencing of the oldest studies (Say-Sallaz et al. 2019), the first half of the twentieth century and before was likely less well covered than the more recent period in our literature review. We added relevant articles from our personal bibliography that did not come up in the systematic search (n = 76 articles, see Randles and Finnegan 2023). This complementary approach corresponded to a long-term, careful exploration and detailed read of articles revolving around the phenology of births in large herbivores. We could identify a substantial number of articles that had not been retrieved by the WoS systematic search, particularly articles published before 1993, those not available online, and those referring to birth phenology in which this topic was not the main focus of the study. In the latter case, birth phenology was not necessarily mentioned in the title, which explains why these articles were overlooked in the systematic search.

We conducted a final selection based on the reading of 145 full articles. From this final screening, we only included the articles directly exploring the phenology of births in large herbivores in wild conditions and their potential drivers (i.e., excluding purely descriptive articles). Finally, our literature review included a total of 124 articles (WoS = 63, personal bibliography = 61, Supplementary Material 2).

### Data extraction from the articles

We extracted manually the following information from each article included in the review: references of the article (e.g., year of publication and journal), references of each population studied (e.g., species vernacular and scientific names, sample size), references of the study site and study period (e.g., country and duration of the study; see Supplementary Material 1 for a detailed description of each variable extracted). Each variable explored in a given population was recorded as one unique observation, which means that an article could contain more than one observation even if a single hypothesis was tested. For instance, an article studying the effect of vegetation phenology on two different populations of caribou consisted of two observations.

For each variable, we extracted the citation of the hypothesis when available (i.e., the formal description of the expectations as formulated by the authors, usually in the Introduction section of the article). In several instances, a variable was tested while the authors did not explicitly state their expectations. For example, we encountered this situation when the authors tested the effect of the sex of the offspring on the timing of birth as a control variable, but did not initially mention if they expected an effect or not, or the direction of the effect. We then classified each variable tested according to 39 factors (e.g., female age, latitude, population density; see Results section for the detailed list), themselves grouped into seven main themes (e.g., the factors mentioned above belong to the themes: female characteristics, environmental characteristics and population characteristics, respectively; see Results section for the detailed list). We identified the parameter of phenology addressed by the variable, i.e., the timing or synchrony of births. We used the codes “supported”, “partially supported” or “unsupported” to report on the outcome of the study. We considered the effect of a variable as “supported” if the associated significance value reported by the authors was below the level alpha = 0.05 (statistical approach). In the absence of a formal statistical test (descriptive approach), we followed the conclusions of the authors and considered the effect of a variable as “supported” when the authors clearly stated so, which could be reported either in the Results or in the Discussion section. In the case of a significant effect detected only on a subset of the population (e.g., when considering male but not female offsprings, or when considering multiparous but not primiparous females), we recorded that the hypothesis was “partially supported”.

### Quality index

To assess the reliability of the hypothesis testing procedure in the articles retrieved for this review, we calculated a general quality index based on the data collection method, the sample size, the study period length and the type of test (statistical or descriptive) used in an article. For each variable tested in an article, we attributed a value between 1 (lowest quality) and 3 (highest quality) to the data collection method, the sample size, the study period length and the type of test conducted (see Supplementary Material 3 for details). We then summed these four values to obtain an index ranging between 4 and 12 for each variable tested. We could not assign a quality index when sample size and/or study period length were not mentioned in the article (n = 26 articles).

To evaluate the empirical support for each theme, we calculated the percentage of variables receiving support (full or partial) versus no support. As our quality index highlighted discrepancies between studies (see Results), we also calculated a weighted mean to account for study reliability. Variables were scored as follows: supported = 1, partially supported = 0.5, unsupported = 0. Weights were assigned based on quality index: no quality index assigned = 1 (n = 112), quality index ranging between 4 and 6 = 2 (n = 6), quality index ranging between 7 and 9 = 3 (n = 167), and quality index ranging between 10 and 12 = 4 (n = 285). A theme was considered to influence timing or synchrony if its weighted mean exceeded 0.5.

Additionally, we explored the change in quality across time using a linear regression with a quadratic effect (assuming a potential stabilisation of quality over time) to model the variation of the quality index of the articles (mean of the quality indexes of all the variables tested in the article) according to the year of publication. We also explored the variation of quality according to the continent and taxonomic family of the populations studied in the articles using ANOVAs. Due to small sample sizes for some taxonomic families (sample size for Antilocapridae, Elephantidae, Hippopotamidae, Moschidae, Rhinocerotidae and Tayassuidae ≤ 3), we grouped Tayassuidae with Suidae, Moschidae with Cervidae, Antilocapridae with Bovidae, and finally, Elephantidae and Hippopotamidae with Rhinocerotidae. All statistical analyses were conducted using the R software (version 4.5.2, R Core Development Team 2025). The data used in the analyses is available from the Zenodo Repository (Anonymised reference).

## Results

### Temporal, geographical and taxonomic focus

We identified a total of 124 articles representing 81 species distributed across 29 countries. The articles were published between 1962 and 2023 in 52 different peer reviewed journals (Fig. 1a), the number of articles increasing quadratically since the 1960’s (Fig. 1b). One article did not directly mention the species names (English et al. 2012) and two articles did not directly mention the country of origin of the populations included in the study (English et al. 2012, Owen-Smith and Ogutu 2013). Almost half (46%) of the species were studied in only one article (e.g., the chital (*Axis axis*), Ahrestani et al. 2011), and a very limited number of species appeared in more than five articles (n = 16 species, e.g., the roe deer appeared in 13 articles, the reindeer and bighorn sheep (*Ovis canadensis*) in 12 articles, Fig. 2). Several populations characterised by continuous long-term monitoring were subject to multiple publications (e.g., the red deer (*Cervus elaphus*) of Rum Island, Scotland; the roe deer of Trois Fontaines, France; the guild of large herbivores of Serengeti National Park, Tanzania). Although the articles retrieved studied between 1 and 80 populations, most focused on one or two populations (n = 96, from the same or different species), and only 11 included more than 10 populations. All the articles including more than 25 populations were review articles which compiled datasets from several original articles to conduct a new set of analyses in a multi-specific or multi-population framework (e.g., English et al. 2012). For the articles providing the duration of the study or the starting and ending dates of data collection for the populations studied (n = 169 populations), the average study length (± standard deviation) was 11 ± 11 years (median [Q1; Q3] = 6 [3; 14] years, Fig. 3).

**Figure 1:**
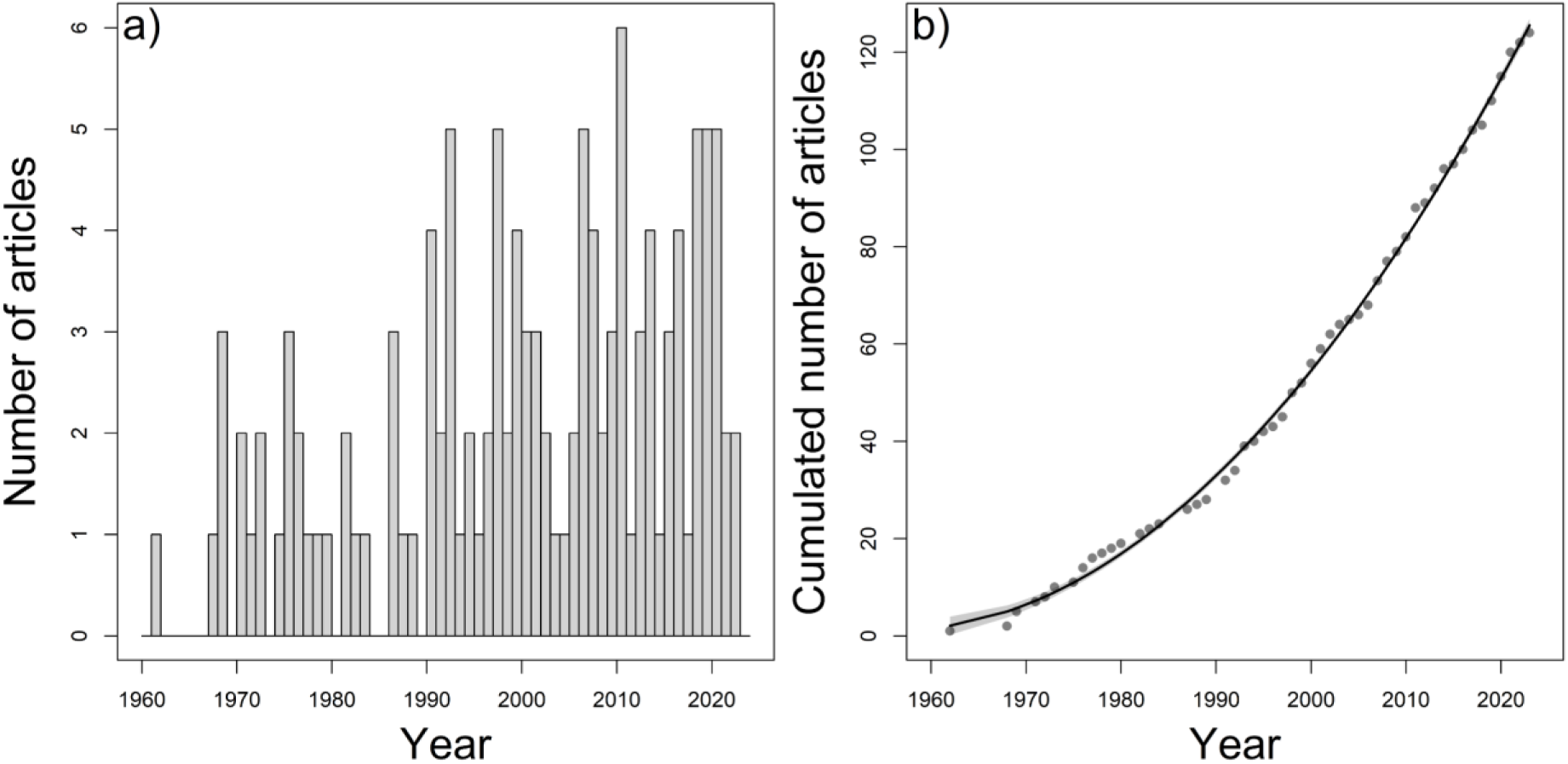
number of articles investigating the drivers of the phenology of birth in large herbivores (n = 124): a) published every year between 1960 and 2024, b) cumulated number of articles published every year (dots: cumulated numbers, line: predictions of the quadratic regression, shaded area: 95% confidence interval).

**Figure 2:**
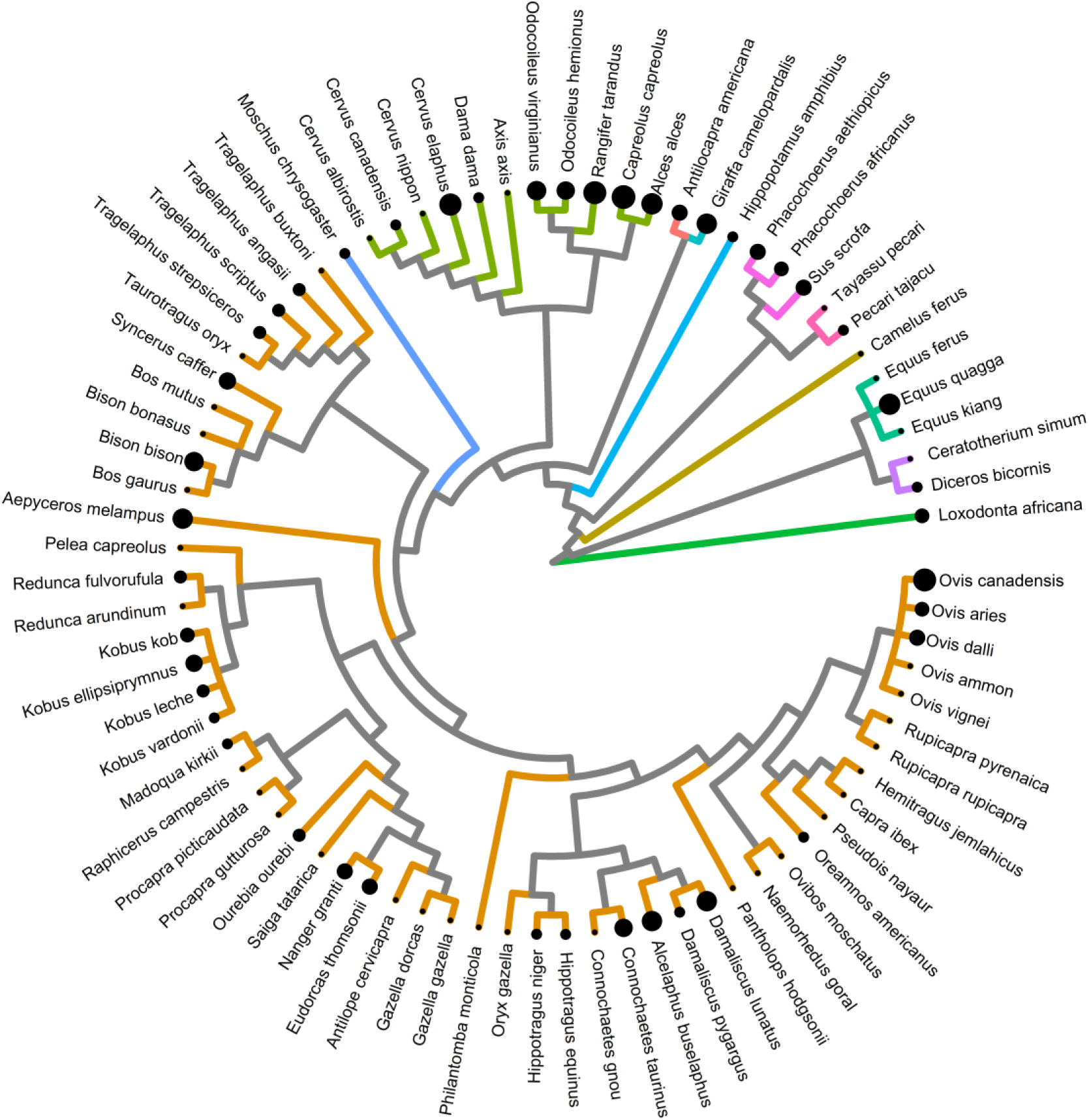
Species studied in the articles investigating the drivers of the phenology of birth in large herbivores (n = 81). The size of the dots is proportional to the number of articles studying a given species (*Ovis aries*, *Ovis gmelini* and *Ovis musimon* were grouped together under *Ovis aries*). The colour of the branches corresponds to taxonomic families: Antilocapridae, Bovidae, Camelidae, Cervidae, Elephantidae, Equidae, Giraffidae, Hippopotamidae, Moschidae, Rhinocerotidae, Suidae, Tayassuidae.

**Figure 3:**
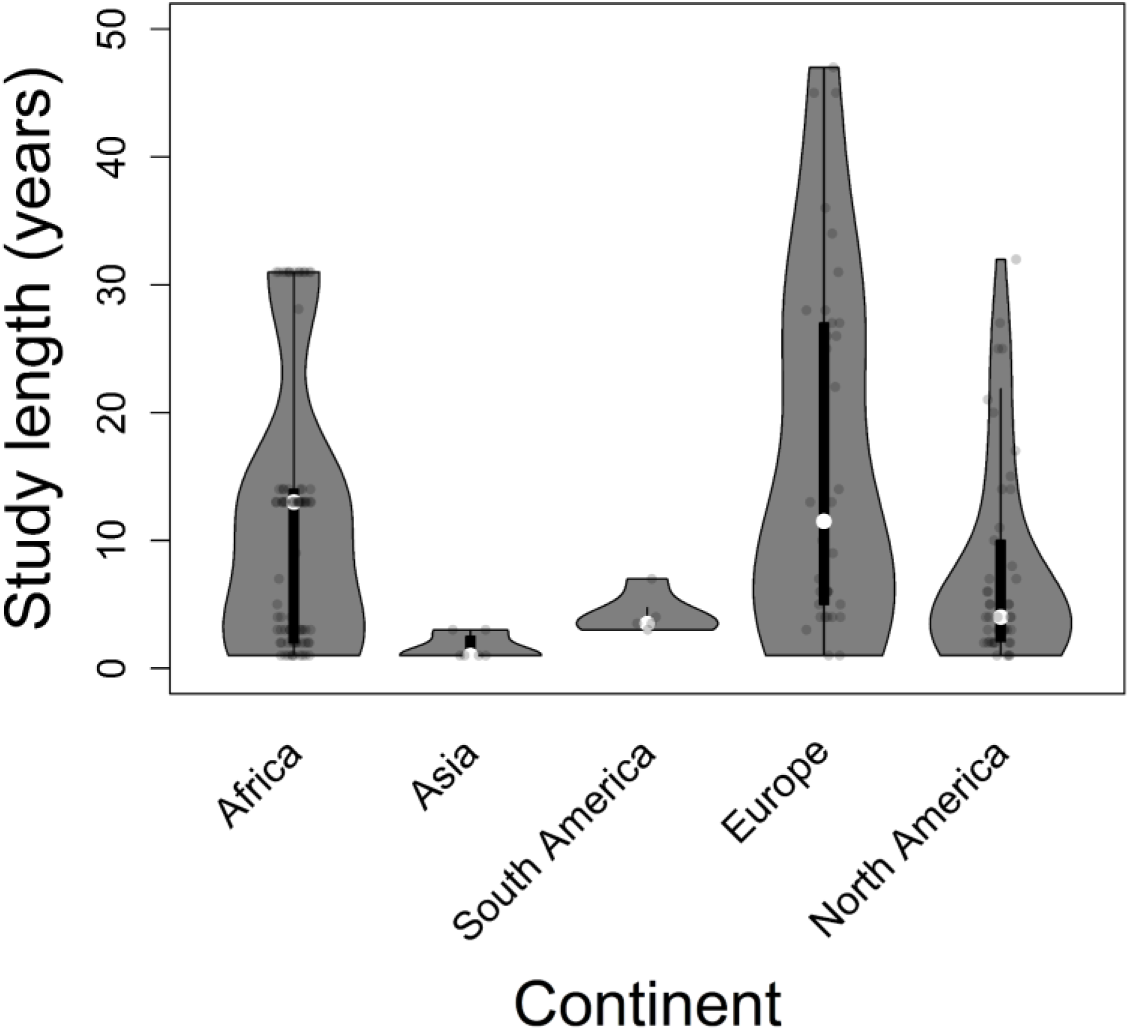
Duration of the studies presented in the articles investigating the drivers of the phenology of births in large herbivores between 1960 and 2024 (n = 169 populations) per continent (in the figure, “South America” stands for “Central and South America”).

North America was the most represented continent with 46 articles on birth phenology, compared to only four from Central and South America (Fig. 4), and three articles covered multiple continents (“worldwide” category in Table 1). The two predominant data collection methods, neonate captures and direct observations, were mainly used in North America and Europe, with the latter also common in Africa (Fig. 5). North American studies additionally relied on movement tracking devices (i.e., identification of parturition date from the mother’s movements and/or identification of juvenile survival from the juvenile’s movements, Supplementary Material 3), and less frequently on vaginal implant transmitters, whereas fetal aging was more common in Africa. Four additional methods were used infrequently: aerial surveys, camera trapping, faecal hormonal assays, and aging of culled individuals.

**Figure 4:**
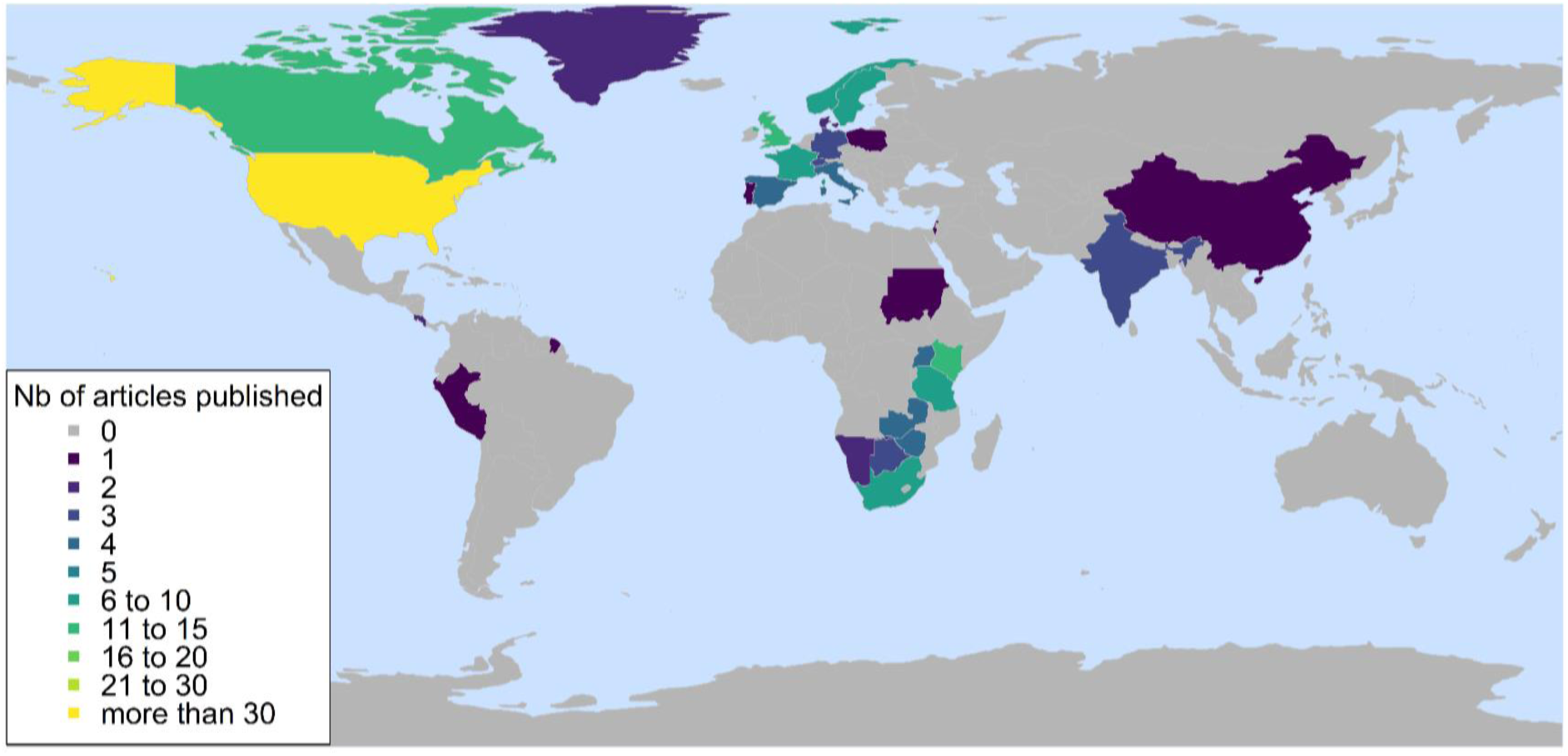
Distribution across countries of the total number of articles investigating the drivers of the phenology of births in large herbivores (n = 124). Some articles cover several countries.

**Figure 5:**
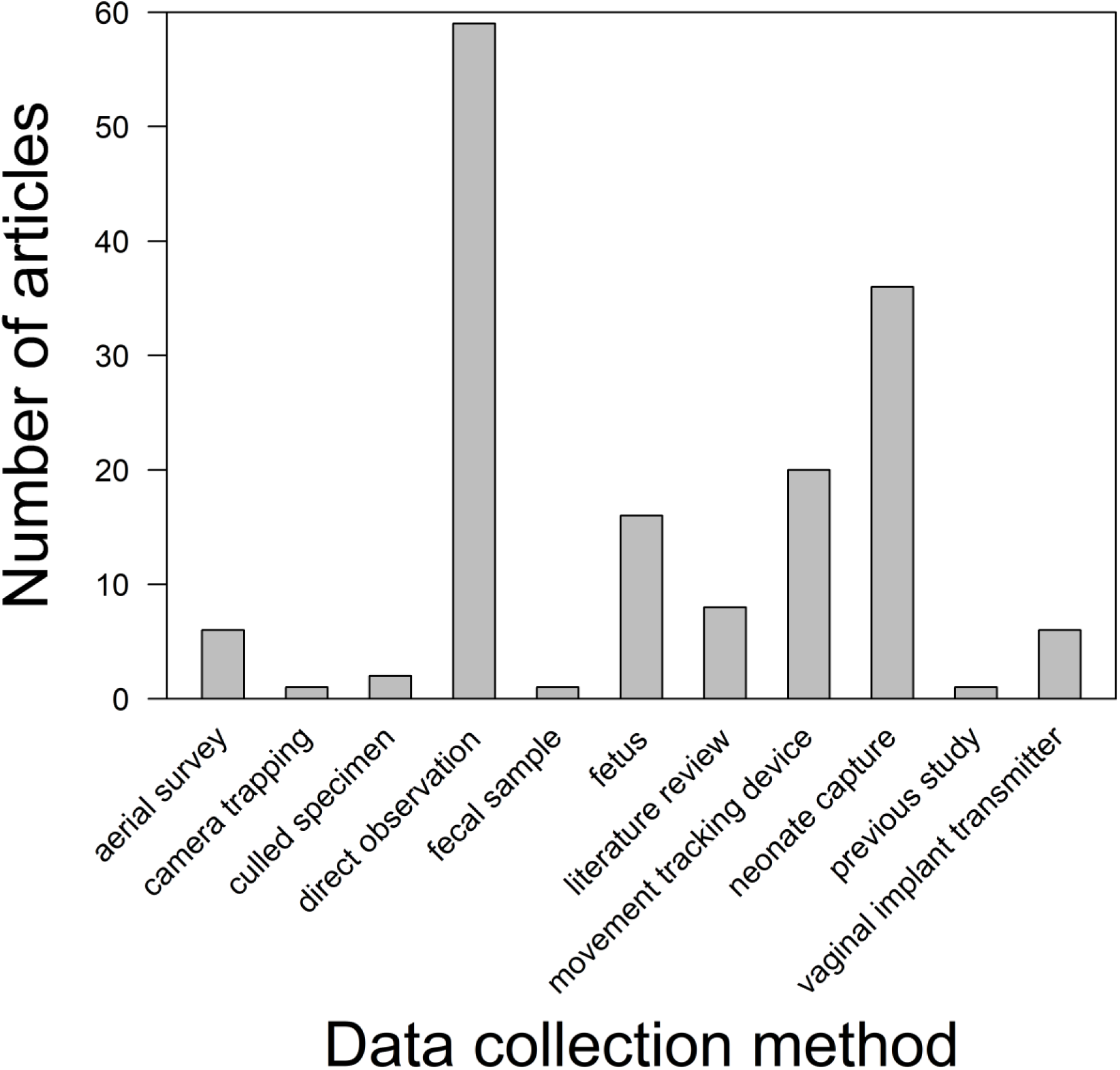
Data collection methods used in the articles investigating the drivers of the phenology of births in large herbivores between 1960 and 2024 (n = 170 populations).

**Table 1:** Number of articles per continent investigating the effect of at least one variable on the timing or on the synchrony of births in large herbivores (the same article can appear in both). The “worldwide” category represents articles studying populations from more than one continent.

| **Continent** | **Number of articles exploring the drivers of the timing of births** | **Number of articles exploring the drivers of the synchrony of births** | **Total number of articles** |
| --- | --- | --- | --- |
| Africa | 35 | 9 | 36 |
| Asia | 4 | 3 | 5 |
| Central and South America | 4 | 0 | 4 |
| Europe | 27 | 8 | 30 |
| North America | 39 | 24 | 46 |
| Worldwide | 1 | 3 | 3 |

### Hypotheses and analytical approach

Over all the recorded variables (n = 570), we identified three main ways to approach the analyses in the articles retrieved. The most common one (n = 311) corresponded to what we defined here as a “closed hypothesis”. In this case, the expectation was clearly stated by the authors (e.g., “*As grass is more seasonal than browse, grazers are expected to have a shorter birth season length than mixed feeders or browsers*”, English et al. 2012). In what we defined as an “open hypothesis” approach (n = 161), an hypothesis was stated by the authors, but there was no specific expectation regarding the effect of the variable tested (e.g., “*We tested relationships between the timing of parturition and maternal age*”, Adams and Dale 1998). In some instances, we could not identify any statement announcing that the effect of a given variable would be tested even though the effect or absence of effect of this variable was reported in the Results section (n = 98). We recorded these cases as the “absence of clear hypothesis” (e.g., Berger 1992 mentioned their hypotheses and expectations after presenting their results: “*Hence the data support the hypothesis that (1) late-breeding females in good body condition shorten gestation, synchronizing births with other females*”). In most cases, the effect of a variable was confirmed by a statistical test (n = 389) rather than a descriptive approach (n = 181). In descriptive approaches, authors tended to use empirical evidence such as a high number of births during the rainy season and not during the rest of the year instead of a statistical test to draw their conclusions. Sixty-one percent of the variables were tested descriptively rather than statistically in articles published on African populations, reaching 67%, on Central and South American populations.

### Quality index

There was a significant relationship between the year of publication and quality index of the articles (n = 103 articles included in the analysis; β_year_ = 5.28 ± 2.25, p = 0.02; β_year*year_ = - 1.31 ± 0.00, p = 0.02; adj. R^2^ = 0.27), with quality increasing over time but progressively reaching a plateau after 2005 (Supplementary Material 3). The quality index varied depending on the continent of the study (F = 13.00, df = 4, p < 0.01), and was significantly larger in Europe and North America than in Africa, Asia and Central and South America (mean difference = 2.06, all p < 0.05; Supplementary Material 3). The quality index also varied depending on the taxonomic family of the population studied (n = 42 species included in the analysis; F = 10.57, df = 5, p < 0.01; Supplementary Material 3). It was significantly larger for Cervidae-Moschidae than for any other taxonomic family (mean difference = 1.73, all p < 0.05), except Girafidae (difference = 1.41, p = 0.07). All other taxonomic families did not show significant differences (all p > 0.05).

### Variables tested

We identified seven themes: i) environmental characteristics, ii) plant characteristics, iii) life history traits, iv) female characteristics, v) offspring characteristics, vi) population characteristics and vii) predation exposure. In each theme we found articles focusing on the timing but also the synchrony of births (Table 2). In some themes, articles were however mainly focused on one or the other (e.g., female, offspring and population characteristics often tested for their effect on timing of births; predation exposure mostly tested for its effect on synchrony of births). These themes encompassed 39 factors (Table 3), themselves encompassing numerous variables. For instance, the factor “rainfall” was investigated via variables such as “monthly rainfall”, “cumulative rainfall”, “total rainfall during the wet season”, “autumn rainfall”, “onset of the rains”. In total, we identified 408 variables whose effect on the timing of birth was tested, whereas we identified 162 variables whose effect was tested on the synchrony of births.

**Table 2:** Number of articles investigating at least once the effect of a factor belonging to one of the seven themes identified in the study on the timing or on the synchrony of births in large herbivores (n = 124).

| **Theme** | **Timing** | **Synchrony** |
| --- | --- | --- |
| Environmental characteristics | 79 | 23 |
| Plant characteristics | 33 | 10 |
| Life history traits | 9 | 9 |
| Female characteristics | 32 | 3 |
| Offspring characteristics | 18 | 1 |
| Population characteristics | 18 | 6 |
| Predation exposure | 6 | 16 |

**Table 3:** Factors (n = 39) tested for their effect on the timing and/or on the synchrony of births in large herbivores, and their corresponding theme (n = 7) as identified in the stu**dy.**

| **Theme** | **Factor** |
| --- | --- |
| Environmental characteristics | altitude, annual climatic seasonality, cloud, day length, inter-annual climatic variability, latitude, local environmental conditions, rainfall, snow, temperature |
| Plant characteristics | faecal or forage protein content, forage biomass, forage quality, general forage quality, general plant phenology, NDVI, plant evapotranspiration |
| Life history traits | anti-predator behaviour of the juvenile, body size, diet, gregariousness, heritability and plasticity, phylogeny |
| Female characteristics | female age, female body mass, female condition, female identity, female migratory strategies, female previous reproductive status, female quality, female social rank |
| Offspring characteristics | offspring inbreeding level, offspring number, offspring sex |
| Population characteristics | population age structure, population density, population sanitary state, population sex-ratio |
| Predation exposure | predation exposure |

When investigating the timing of births, rainfall was the most common factor tested (“environmental characteristics” theme, in n = 28 articles). Female age was the most investigated factor among female characteristics (in n = 19 articles), while it was plant phenology in plant characteristics (in n = 14 articles), offspring sex in offspring characteristics (in n = 13 articles) and population density in population characteristics (in n = 12 articles, Supplementary Material 4). When investigating the synchrony of births, predation exposure was the most common factor tested (in n = 16 articles). Latitude was the most investigated factor among environmental characteristics (in n = 9 articles), while it was the anti-predator behaviour of the juvenile in the “life history traits” theme (in n = 7 articles).

The effects of plant characteristics as well as environmental characteristics were commonly explored in all continents (Fig. 6). In Africa, they were mainly explored via rainfall and annual climatic seasonality (mostly based on a broad description of the temporal succession of wet and dry seasons), while these themes were often described via local environmental conditions (i.e., difference of seasonality and climate based on spatial differences) in Europe, and plant phenology as well as temperature in North America. In addition, the effects of female and population characteristics were also commonly explored in Europe and North America, mostly via female quality and identity as well as population density in Europe, and via female age and previous reproductive status in North America. The effects of predation exposure as well as offspring characteristics were also commonly explored in North America. When considering the effect of a variable on the timing of births, 50% or more tests received partial or full support in all themes, except from the theme “offspring characteristics”, where, to the contrary, more than 75% received no support (Fig. 7a). When accounting for the quality of the tests, the weighted means revealed sensibly similar results than unweighted means, despite the fact that plant and population characteristics received slightly less than 50% support (Table 4). When considering the effect of a variable on the synchrony of births, life history traits, female characteristics and offspring characteristics (but n = 1 in the latter) received largely more than 50% support, even after accounting for the quality of the tests (Fig. 7b, Table 4). The other themes received between 25 and 50% support overall (see details in Supplementary Material 5).

**Figure 6:**
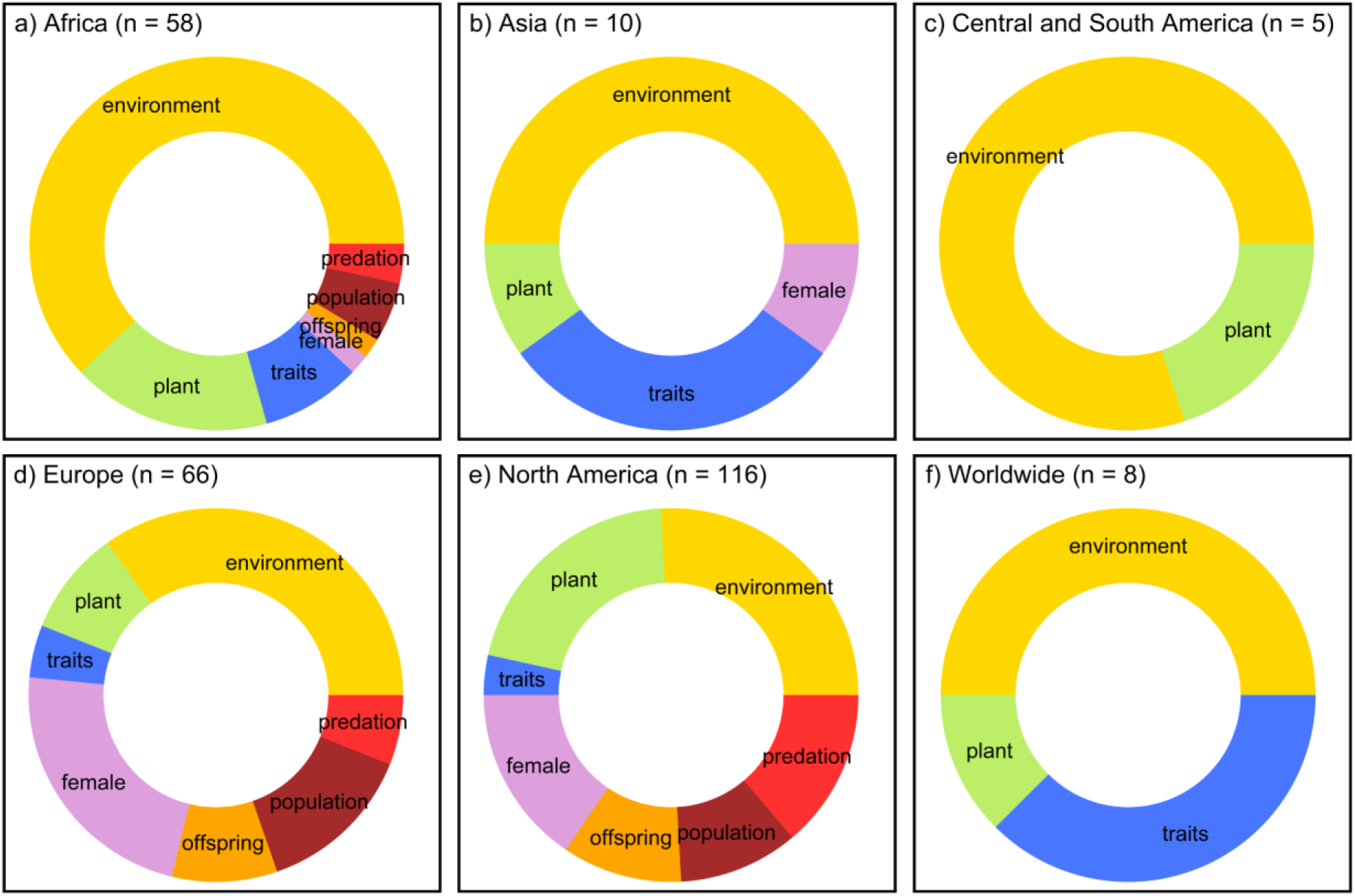
Frequency of investigation of each theme (based on the number of articles exploring at least once a factor of the theme considered) in the articles investigating the drivers of the phenology of births in large herbivores between 1960 and 2024: environmental characteristics; plant characteristics; life history traits; female characteristics; offspring characteristics; population characteristics; predation exposure.

**Figure 7:**
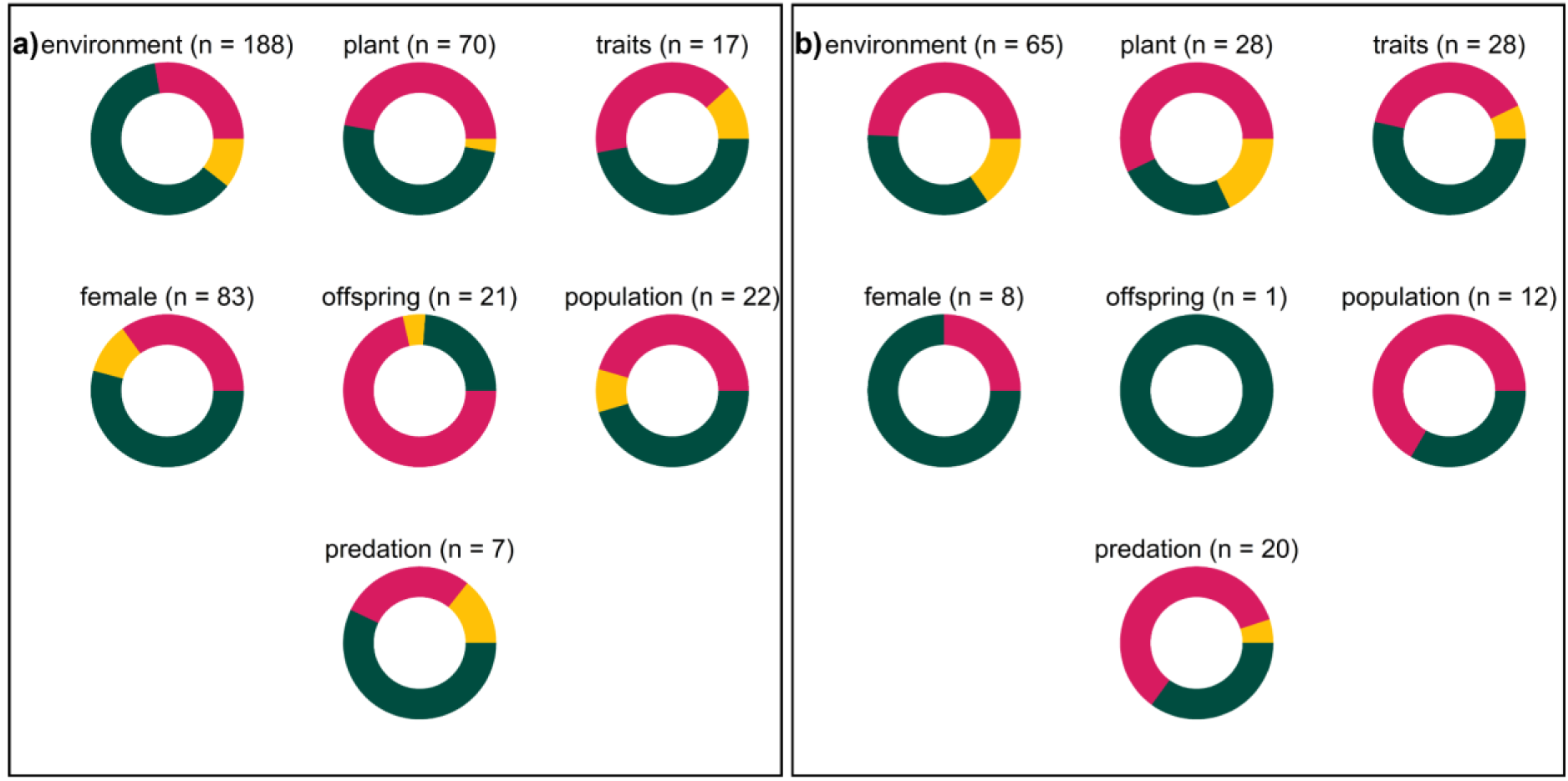
Proportion of tests receiving support (green), partial support (orange) or no support (red) in the articles investigating the drivers of the phenology of birth in large herbivores: a) timing and b) synchrony, for each theme (n = 7; environmental characteristics; plant characteristics; life history traits; female characteristics; offspring characteristics; population characteristics; predation exposure).

**Table 4:** Weighted means illustrating the strength of support received by each theme (environmental characteristics; plant characteristics; life history traits; female characteristics; offspring characteristics; population characteristics; predation exposure) when investigating the drivers of the phenology of birth in large herbivores (timing and synchrony), while accounting for the quality of the test (n = 570, theme specific sample sizes reported in the table). We considered that a given theme received support if the corresponding weighted mean was > 0.5.

| Theme | Timing | Synchrony |
| --- | --- | --- |
| Environmental characteristics | 0.70 ($n = 188$) | 0.49 ($n = 65$) |
| Plant characteristics | 0.48 ($n = 70$) | 0.34 ($n = 28$) |
| Life history traits | 0.57 ($n = 17$) | 0.61 ($n = 28$) |
| Female characteristics | 0.58 ($n = 83$) | 0.71 ($n = 8$) |
| Offspring characteristics | 0.24 ($n = 21$) | 1.00 ($n = 1$) |
| Population characteristics | 0.48 ($n = 22$) | 0.34 ($n = 12$) |
| Predation exposure | 0.67 ($n = 7$) | 0.37 ($n = 20$) |

## Discussion

### Support for the dominant hypotheses: seasonality and predation

Even after controlling for the quality of the tests, more than half of the tests exploring the effect of an environmental characteristic and around half of the tests exploring the effect of a plant characteristic concluded for the existence of an effect on the timing of births (e.g., Bunnell 1982, Sinclair et al. 2000, Moyes et al. 2011, Renaud et al. 2022) from a variety of approaches and variables. Almost 20 factors were used (e.g., rainfall, temperature, greenness indexes, forage biomass, plant evapotranspiration). The effect of rainfall was mainly studied in Africa, and the one of temperature mainly in the northern hemisphere (Europe and North America), in accordance with the region-specific limiting factor of vegetation growth, the southern hemisphere seasonality being predominantly determined by rainfall (wet versus dry seasons) and the northern hemisphere seasonality by temperature (hot versus cold seasons, Owen-Smith and Ogutu 2013). The tests exploring the effect of an environmental or a plant-related variable found less support for the existence of an effect on synchrony (48% and 34% respectively, after controlling for the quality of the tests, Table 4) than timing of births. These less conclusive findings may result from a complex interaction between multiple drivers, to which synchrony may be more sensitive than timing, such as social factors and exposure to predation (Gosling 1969).

The articles retrieved in the present literature review explored the effect of predation pressure predominantly via two approaches, the direct impact of predation on juvenile mortality and the anti-predator behaviour of the juvenile. They found moderate support (e.g., Michel et al. 2020), as only 40% of the tests supported or partially supported the effect of predation on synchrony (37% when controlling for the quality of the tests, Table 4), and 71% of the tests supported or partially supported the effect of predation on individual birth timing (67% when controlling for the quality of the tests, Table 4). The articles using the anti-predator behaviour of the juvenile were often descriptive (e.g., Ogutu et al. 2013, but see Rutberg 1987 for a statistical approach), which can affect the strength of the conclusions. Additionally, the anti-predator behaviour of the juvenile was generally classified according to the hider/follower dichotomy (Lent 1974). However, this trait correlates with other life history traits such as group size and sociality, diet and body size of the species, which might themselves affect birth phenology (Sinclair et al. 2000). The classification of certain species at one end or the other of the gradient is therefore debated (e.g., African buffalo (*Syncerus caffer*) and plains bison (*Bison bison*), Green and Rothstein 1993b, Ryan et al. 2007), making it more difficult to establish general and universal conclusions based on this approach. From the present literature, we conclude that predation pressure might have a significant effect on the synchrony of births in African species (effect supported in 63% of the tests, 58% after controlling for the quality of the tests). African ecosystems are less subject to food resource and climate seasonality than, for instance, northern temperate ecosystems, in which predation seems to have less effect than climate or vegetation phenology on the distribution of births (effect partially or fully supported in only 32% of the tests, 34% after controlling for the quality of the tests). Such a finding corroborates the results of previous comparative studies (Zerbe et al. 2012).

Note that some themes were tested several times in the same article (e.g. Sinclair et al. (2000) accounted for 24 observations for the effect of plant characteristics on birth timing), and some species or populations appeared multiple times in our dataset (e.g. the roe deer accounted for 15 observations and, more particularly, the roe deer from Trois Fontaines, France accounted for 4 observations when testing the effect of environmental characteristics on birth timing, Supplementary Material 6). By treating them as independent observations when quantifying the support received by each theme, our approach may have inflated the influence of these well-studied systems on our results.

### Emerging research theme avenues

Although fairly frequently explored in European and North American populations, the effects of female, offspring and population characteristics were marginally accounted for in the study of the phenology of births in large herbivores. When investigated, evidence of their effect is mixed. It remains unclear if this lack of consistency in the results comes from biological reasons, these drivers potentially acting as a secondary driver after environmental conditions and mostly at the individual level might have a limited effect on the phenology of births, or from methodological reasons. The investigation of such drivers requires a detailed understanding in the long-term of the population studied and individually-centred monitoring, which can be challenging to implement and is thus infrequently done or might lead to imperfect datasets (e.g., data collected for a small sample of the population or biased toward certain individuals, data collected on a short period of time, Festa-Bianchet et al. 2017). Accordingly, we found that the articles focusing on European and North American populations (more particularly Cervidae populations with long-term monitoring) extensively exploring these drivers were also characterised by higher quality indexes than the other articles. Due to the difficulty to collect accurate data as well as their expected secondary role in the determination of the timing of births, these drivers are often included as control or secondary variables. Although these characteristics are often related to environmental conditions (e.g., the reproductive status of the female during the previous reproductive season can be related to forage availability and climatic conditions), intrinsic characteristics such as female age, social rank and experience (e.g., primiparity or multiparity) have also been identified as relevant drivers of the phenology of births (e.g., Berger and Cain 1999, Bonnet et al. 2019). Even if these variables do not actively influence the phenology of births, they nevertheless mechanically affect the distribution (e.g., in a population of mainly young females, one can expect the timing of births to be delayed), and failing to take these demographic processes into account can affect conclusions when evolutionary or plastic drivers such as environmental seasonality are explored. Consequently, population structure should not be disregarded when studying the phenology of births.

Besides, despite directly affecting the phenology of birth, food resources and environmental characteristics can also affect female’s ability to breed, the timing of breeding, and finally the timing of parturition (Hogg et al. 2017). However, articles dealing exclusively with the phenology of mating were not considered in this review (e.g., Dauphiné and McClure 1974). Gestation length is adjustable within some limits (Clements et al. 2011, Zerbe et al. 2012), so the phenology of mating cannot be regarded as an absolute reflection of the phenology of birth. Additionally, mating is more difficult to observe in the field than birth and does not necessarily lead to a birth (i.e., abortion). We trust the number of articles in this field is limited and their exclusion should not affect the general conclusions of our study. Finally, the effect of diseases (e.g., Berger and Cain 1999) and inbreeding (e.g., Bonnet et al. 2019) prevalence on the phenology of births is currently under-explored. Several pathogens can cause abortion in large herbivores (e.g., brucellosis (*Brucella abortus*), Cross et al. 2015), thus leading to the possibility for an altered phenology of births in infected populations, depending on the prevalence of the disease in the population and the timing of abortion. Similarly under-explored, human interventions such as hunting (e.g., Gamelon et al. 2011) or active management (Kaze et al. 2016) might affect the evolutionary and/or plastic determination of the synchrony of births in a population. Gamelon et al. (2011) showed that hunting acts as a selection pressure for earlier dates of birth in a population of wild boar (*Sus scrofa*), as being born earlier increases development time and allows females to reproduce earlier, an evolutionary advantage when mortality is high. Kaze et al. (2016) hypothesised that a better management of bison natural habitat via environmental rehabilitation and reduction of competition with livestock for resource access enhanced female condition and increased birth synchrony. In the current context of climate change, modification of land use and development of human activities across the globe, there is a need for a better understanding of the effect of such anthropogenic and epidemiological factors on the phenology of births in large herbivores, and more broadly in mammals.

### Research challenges

As illustrated in this study, the phenology of births is driven by a combination of drivers, which might be difficult to separate and describe. Although most of the articles used “direct tests”, when the variable of interest is directly used in the test (e.g., effect of the age of the female on the date of parturition, Bon et al. 1993), the use of “tests via proxy” was common (when the variable of interest is not available or difficult to measure in the field leading to the use of a proxy instead, e.g., effect of food availability via the variability of rainfall, Ndibalema 2009). Additionally, some articles also used “logical tests”, when the variable of interest is not available or difficult to measure in the field, and a theoretical reasoning is used to draw conclusions from the observations (e.g., “*if synchrony in births is primarily an antipredator adaptation by ungulates, then episodic and unpredictable droughts and floods should not significantly shift the timing and synchrony of births in seasonally breeding ungulates*”, Ogutu et al. 2010). However, in such cases, the effect of the driver tested cannot be proven conclusively.

The importance of interacting and cascading effects is frequently highlighted in the Discussion section or even tested in the articles retrieved for the present literature review (e.g., Stopher et al. 2008, Zerbe et al. 2012, Aikens et al. 2021). The variability of our findings in terms of the support received by each factor tested also illustrates this complexity. To address this complexity, the use of aggregated indices (i.e., one index combining several variables) is frequent, such as: i) latitude which combines factors such as temperature, day length, vegetation type, seasonality of the environment (e.g., English et al. 2012), ii) anti-predator behaviour of the juvenile which refers to the level of precocity of the juvenile but also correlates with other life history traits (e.g., Sinclair et al. 2000), iii) site-specific comparisons which uses spatial variability to describe different levels of exposure to predation for example, but also includes variable local environmental conditions (e.g., Post et al. 2003). Such an approach can provide a general understanding of a set of factors on the phenology of births, but limits our understanding of individual effects. Although the possibilities are limited when studying populations in natural conditions (Clauss et al. 2021), the study of the phenology of births in systems where one or several factors are constant or fairly stable can help in this regard.

Most of the factors tested in the articles retrieved correspond to proximal factors inducing a short-term response fairly easy to monitor. However, monitoring the effect of ultimate factors generating slow evolutionary responses is more challenging. Such investigations require comparative analyses involving multi-species comparisons controlling for genetic relatedness (Zerbe et al. 2012, Bonnet et al. 2019). The majority of the articles retrieved for the present literature review focused on a single species, and the rare multi-species studies usually used descriptive comparisons between species based on life history traits and/or did not account for phylogenetic correlations (e.g., Gosling 1969, Rutberg 1987). Large scale comparative analyses can be of particular interest to empirically test theoretical hypotheses about life history traits and evolutionary responses in reproductive phenology (Rutberg 1987, Zerbe et al. 2012).

Lastly, our semi-systematic approach (i.e. combination of a systematic search based on keywords using title screening on the Web of Science and a non-systematic search using in depth, manual exploration of the birth phenology literature) revealed that the literature on the phenology of births in large herbivores is currently not easy to identify: only half of the relevant articles retrieved for this study came from the systematic search. Although our semi-systematic approach might reduce repeatability for future similar studies, it illustrates that a screening solely based on the title of the articles would severely underestimate the number of relevant articles found. For articles whose main focus is the phenology of births, we encourage the use of clear titles, and for those where it is a secondary topic, to include words such as “phenology” in the keywords or abstract. This will ensure that relevant articles can be more easily identified and referenced in search engines (Pottier et al. 2024), improving repeatability and comparability between studies exploring the drivers of the phenology of births.

### Geographic and taxonomic gap

Our literature review identified a large geographical and taxonomic gap in the study of the phenology of births in large herbivores, likely to be generalized to other mammals too. More than 90% of the articles retrieved concern Africa, Europe and North America, leaving Asia and Central and South America mostly unexplored. At a finer geographical scale, West and Central Africa as well as Eastern Europe also have almost no data available, or are concerned by purely descriptive studies (e.g., in Honduras, Klein 1982; in Nepal, Bhusal et al. 2020). Most of these ecosystems are characterised by dense vegetation, difficult conditions and/or remote locations from human infrastructures, explaining the overall gap in the scientific literature (Kelt and Meserve 2014). Additionally, the large herbivores occupying these environments are often elusive, mostly nocturnal and/or occurring at low densities (e.g., Cephalophus and Philantomba genera in Central Africa, Zerbe et al. 2012), which makes their study challenging, as illustrated by the lower quality index of these articles.

Moreover, although seasonal in terms of food availability, the absence of major environmental seasonality in equatorial ecosystems (e.g., photoperiod, temperature) might also have played a role in the lack of studies conducted under these latitudes (Bronson 2009), and led to the assumption that no reproductive seasonality should be expected, thus reducing the interest for the exploration of the phenology of births. However, there is evidence for reproductive seasonality around the equator, as illustrated in other taxa such as Chiroptera (e.g., Heideman and Utzurrum 2003), Primates (e.g. Dezeure et al. 2022), and in rare examples in the present literature review (e.g., the phenology of births of the collared peccary (*Pecari tajacu*) is correlated with fruiting phenology in French Guiana, Dubost and Henry 2017). Additionally, the investigation of the phenology of births in species under stable environmental conditions (compared to temperate systems characterised by dramatic seasonal changes) opens the door to the exploration of other drivers such as predator pressure and social characteristics (e.g., in this study, Sinclair et al. 2000; in Chiroptera, Porter and Wilkinson 2001) by providing natural experimental conditions where the dominant driver regulating the phenology of births in large herbivores, namely environmental seasonality, is controlled. Such a framework can help better understand the role of secondary drivers when environmental seasonality is not a strong constraint (Zerbe et al. 2012). Finally, even in the absence of reproductive seasonality, investigating the phenology of births is critical for the understanding of its cascading effects on population dynamics, in particular in the context of climate change (e.g., the increase of winter births due to earlier onset of plant growth is negatively correlated with juvenile recruitment in feral cattle (*Bos taurus*), Burthe et al. 2011).

## Data availability statement

Data available from the Zenodo Repository: https://doi.org/10.5281/zenodo.17483679 (Anonymised reference).

## Authors’ contributions

Lucie Thel: Conceptualization - Methodology - Formal analysis - Investigation - Data Curation - Writing: Original Draft - Writing: Review & Editing - Visualisation - Funding acquisition – Validation

Christophe Bonenfant: Conceptualization - Methodology - Writing: Review & Editing - Visualisation - Supervision - Funding acquisition – Validation Simon Chamaillé-Jammes: Conceptualization - Methodology - Writing: Review & Editing - Supervision - Funding acquisition – Validation

## Conflict of interest statement

The authors declare that they have no conflict of interest.

## Acknowledgments

We thank Elise Huchard for her suggestions during the early-stage conceptualization of the project and Hermanus Swanepoel for his help in data collection. This work was supported by a grant from the “Ministère Français de l’Enseignement Supérieur, de la Recherche et de l’Innovation” through the “Ecole Doctorale E2M2” of the “Université Claude Bernard Lyon 1” as well as Postgraduate Research Scholarship funding from Nelson Mandela University. The authors thank two anonymous reviewers for their insightful comments on a previous draft.

## Notes

### Competing Interest Statement

The authors have declared no competing interest.

### Summary of Updates

Main revision of the introduction and sections of the discussion for publication process.

