## supplementary material for "Evolutionary and ecological determinants of the phenology of births in wild large herbivores, a systematic review"

### Supplementary Material 1: Data collection process for the systematic review

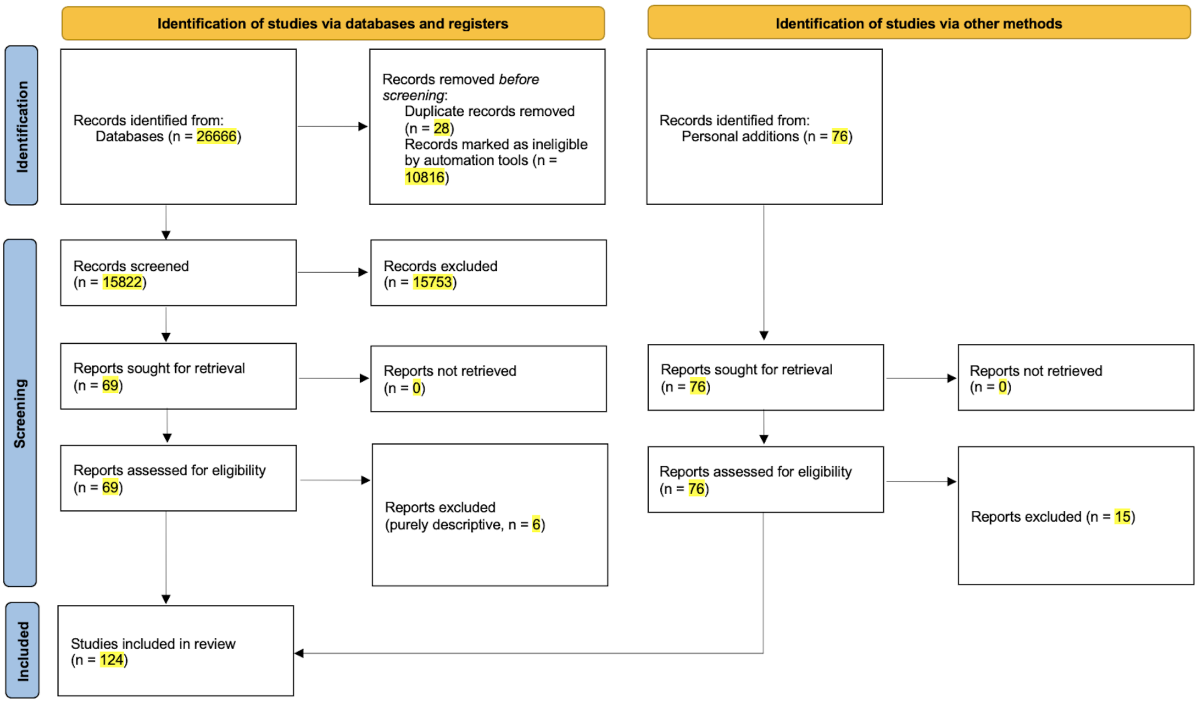


Figure S1.1: PRISMA diagram of the systematic literature review.

Table S1.1: Information extracted from the articles included in the review (n = 124).

| Column name | Column description |
|---|---|
| **paper_ref** | family name of the first author and year of publication |
| **paper_first_author** | family name of the first author |
| **paper_publi_year** | year of publication |
| **paper_title** | full title of the article |
| **paper_journal** | name of the journal which published the article |
| **pop_nb** | number of populations studied (same or different species) |
| **pop_ref** | ID number of the population studied (if only one population studied, pop_ref = 1) |
| **pop_data_coll_method** | how was the information about the dates of birth collected |
| **pop_sample_size** | total number of births recorded for the analysis (alternatively, number of parturitions) |
| **species_vernacular_name** | vernacular name of the species (as used by the authors) |
| **species_latin_name** | scientific name of the species (genus and specific epithet), following the classification of the IUCN Red List, 2024 (https://www.ultimateungulate.com/Taxonomy.html) |
| **study_site_continent** | continent where the study took place |
| **study_site_country** | country (or geographic unit when relevant, e.g., French Guiana) where the study took place |
| **study_site_name** | name of the location of the study site (as used by the authors) |
| **study_period_start** | first year of data collection |
| **study_period_end** | last year of data collection |
| **study_period_length** | number of years for which the data was collected (accounting for missing years when reported by the authors, alternatively number of years elapsed between the first and last year of data collection) |
| **hypothesis_id** | ID number of the variable tested |
| **hypothesis_tested_nb** | number of variables tested in the article |
| **hypothesis_pheno_char** | phenology characteristic the test focused on (i.e., timing or synchrony) |

| Column name | Column description |
|---|---|
| **hypothesis_type** | was the hypothesis tested statistically or descriptively |
| **hypothesis_validated** | was the hypothesis supported, partially supported or unsupported (according to the authors) |
| **test_rationale** | was the hypothesis open, closed or not clearly stated |
| **test_variable** | keywords describing the variable tested |
| **test_factor** | factor describing the variable tested |
| **test_theme** | theme describing the factor tested |
| **paper_doi** | doi of the article |
| **paper_source** | how was the article found (Web of Science search or personal bibliography) |
| **quality_global_index** | quality index defined according to data collection method, sample size, study period length and type of test, as defined in Supplementary Material 3 |

**Supplementary Material 2: References of the articles included in the systematic review (n = 124)**


Adams, L. G. and Dale, B. W. 1998. "Timing and synchrony of parturition in Alaskan caribou." *Journal of Mammalogy* 79 (1): 287-294.

Adams, L. G., Singer, F. J. and Dale, B. W. 1995. "Caribou calf mortality in Denali national park, Alaska." *The Journal of Wildlife Management* 59 (3): 584-594.

Ahrestani, F. S., Van Langevelde, F., Heitkönig, I. M. A. and Prins, H. H. T. 2012. "Contrasting timing of parturition of chital *Axis axis* and gaur *Bos gaurus* in tropical South India–the role of body mass and seasonal forage quality." *Oikos* 121 (8): 1300-1310.

Aikens, E. O., Dwinnell, S. P. H., LaSharr, T. N., Jakopak, R. P., Fralick, G. L., Randall, J., Kaiser, R., Thonhoff, M., Kauffman, M. J. and Monteith, K. L. 2021. "Migration distance and maternal resource allocation determine timing of birth in a large herbivore." *Ecology* 102 (6): e03334.

Allsopp, R. 1971. "Seasonal breeding in bushbuck (*Tragelaphus scriptus* Pallas, 1776)." *African Journal of Ecology* 9 (1): 146-149.

Andersen, R. and Linnell, J. D. 1998. "Ecological correlates of mortality of roe deer fawns in a predator-free environment." *Canadian Journal of Zoology* 76 (7): 1217-1225.

Anderson, J. L. 1979. "Reproductive seasonality of the nyala *Tragelaphus angasi*; the interaction of light, vegetation phenology, feeding style and reproductive physiology." *Mammal Review* 9 (1): 33-46.

Baharav, D. 1983. "Reproductive strategies in female mountain and dorcas gazelles (*Gazella gazella gazella* and *Gazella dorcas*)." *Journal of Zoology* 200 (4): 445-453.

Barber-Meyer, S. M., Mech, L. D. and White, P. J. 2008. "Elk calf survival and mortality following wolf restoration to Yellowstone National Park." *Wildlife Monographs* 169 (1): 1-30.

Berger, J. 1992. "Facilitation of reproductive synchrony by gestation adjustment in gregarious mammals: a new hypothesis." *Ecology* 73 (1): 323-329.

Berger, J. and Cain, S. L. 1999. "Reproductive synchrony in brucellosis-exposed bison in the southern Greater Yellowstone Ecosystem and in noninfected populations." *Conservation Biology* 13 (2): 357-366.

Boertje, R. D., Frye, G. G. and Young Jr, D. D. 2019. "Lifetime, known-age moose reproduction in a nutritionally stressed population." *The Journal of Wildlife Management* 83 (3): 610-626.

Bon, R., Dardaillon, M. and Estevez, I. 1993. "Mating and lambing periods as related to age of female mouflon." *Journal of Mammalogy* 74 (3): 752-757.

Bonnet, T., Morrissey, M. B., Morris, A., Morris, S., Clutton-Brock, T. H., Pemberton, J. M. and Kruuk, L. E. B. 2019. "The role of selection and evolution in changing parturition date in a red deer population." *PLoS Biology* 17 (11): e3000493.

Bowyer, R. T. 1991. "Timing of parturition and lactation in southern mule deer." *Journal of Mammalogy* 72 (1): 138-145.

Bowyer, R. T., Van Ballenberghe, V. and Kie, J. G. 1998. "Timing and synchrony of parturition in Alaskan moose: long-term versus proximal effects of climate." *Journal of Mammalogy* 79 (4): 1332-1344.

Bunnell, F. L. 1980. "Factors controlling lambing period of Dall's sheep." *Canadian Journal of Zoology* 58 (6): 1027-1031.

Bunnell, F. L. 1982. "The lambing period of mountain sheep: synthesis, hypotheses, and tests." *Canadian Journal of Zoology* 60 (1): 1-14.

Cameron, R. D., Smith, W. T., Fancy, S. G., Gerhart, K. L. and White, R. G. 1993. "Calving success of female caribou in relation to body weight." *Canadian Journal of Zoology* 71 (3): 480-486.

Carmichael, I. H., Patterson, I., Drager, N. and Breton, D. A. 1977. "Studies on reproduction in the African buffalo (*Syncerus caffer*) in Botswana." *South African Journal of Wildlife Research* 7 (2): 45-52.

Chen, W., Adamczewski, J. Z., White, L., Croft, B., Gunn, A., Football, A., Leblanc, S. G., Russell, D. E. and Tracz, B. 2018. "Impacts of climate-driven habitat change on the peak calving date of the Bathurst caribou in Arctic Canada." *Polar Biology* 41 (5): 953-967.

Clutton-Brock, T. H., Major, M., Albon, S. D. and Guinness, F. E. 1987. "Early development and population dynamics in red deer. I. Density-dependent effects on juvenile survival." *Journal of Animal Ecology* 56: 53-67.

Côté, S. D. and Festa-Bianchet, M. 2001. "Birthdate, mass and survival in mountain goat kids: effects of maternal characteristics and forage quality." *Oecologia* 127 (2): 230-238.

Couriot, O. H., Cameron, M. D., Joly, K., Adamczewski, J., Campbell, M. W., Davison, T., Gunn, A., Kelly, A. P., Leblond, M., Williams, J., Fagan, W. F., Brose, A. and Gurarie, E. 2023. "Continental synchrony and local responses: Climatic effects on spatiotemporal patterns of calving in a social ungulate." *Ecosphere* 14 (1): e4399.

Dasmann, R. F. and Mossman, A. S. 1962. "Reproduction in some ungulates in Southern Rhodesia." *Journal of Mammalogy* 43 (4): 533-537.

Deutsch, J. C. and Ofezu, A. A. 1994. "Uganda kob reproductive seasonality: optimal calving seasons or condition-dependent oestrus?". *African Journal of Ecology* 32 (4): 283-295.

Dubost, G. and Henry, O. 2017. "Seasonal reproduction in neotropical rainforest mammals." *Zoological studies* 56 (2): e2.

English, A. K., Chauvenet, A. L. M., Safi, K. and Pettorelli, N. 2012. "Reassessing the determinants of breeding synchrony in ungulates." *PLoS One* 7 (7): e41444.

Estes, R. D. 1976. "The significance of breeding synchrony in the wildebeest." *African Journal of Ecology* 14 (2): 135-152.

Fairall, N. 1968. "The reproductive seasons of some mammals in the Kruger National Park." *African Zoology* 3 (2): 189-210.

Fairbanks, W. S. 1993. "Birthdate, birthweight, and survival in pronghorn fawns." *Journal of Mammalogy* 74 (1): 129-135.

Feder, C., Martin, J. G. A., Festa-Bianchet, M., Bérubé, C. and Jorgenson, J. 2008. "Never too late? Consequences of late birthdate for mass and survival of bighorn lambs." *Oecologia* 156 (4): 773-781.

Festa-Bianchet, M. 1988. "Birthdate and survival in bighorn lambs (*Ovis canadensis*)." *Journal of Zoology* 214 (4): 653-661.

Field, C. R. and Blankenship, L. H. 1973. "Nutrition and reproduction of Grant's and Thomson's gazelles, Coke's hartebeest and giraffe in Kenya." *Journal of Reproduction and Fertility* 19: 287-301.

Flydal, K. and Reimers, E. 2002. "Relationship between calving time and physical condition in three wild reindeer *Rangifer tarandus* populations in southern Norway." *Wildlife Biology* 8 (2): 145-151.

Freeman, E. D., Larsen, R. T., Peterson, M. E., Anderson Jr, C. R., Hersey, K. R. and Mcmillan, B. R. 2014. "Effects of male-biased harvest on mule deer: implications for rates of pregnancy, synchrony, and timing of parturition." *Wildlife Society Bulletin* 38 (4): 806-811.

Freeman, E. W., Whyte, I. and Brown, J. L. 2009. "Reproductive evaluation of elephants culled in Kruger National Park, South Africa between 1975 and 1995." *African Journal of Ecology* 47 (2): 192-201.

Froy, H., Martin, J., Stopher, K. V., Morris, A., Morris, S., Clutton-Brock, T. H., Pemberton, J. M. and Kruuk, L. E. B. 2019. "Consistent within-individual plasticity is sufficient to explain temperature responses in red deer reproductive traits." *Journal of Evolutionary Biology* 32 (11): 1194-1206.

Fryxell, J. M. 1987. "Seasonal reproduction of white-eared kob in Boma National Park, Sudan." *African Journal of Ecology* 25 (2): 117-124.

Fuller, T. K., Silva, A. M., Montalvo, V. H., Sáenz-Bolaños, C. and Carrillo, J. E. 2020. "Reproduction of white-tailed deer in a seasonally dry tropical forest of Costa Rica: a test of aseasonality." *Journal of Mammalogy* 101 (1): 241-247.

Gaillard, J. M., Delorme, D., Jullien, J. M. and Tatin, D. 1993. "Timing and synchrony of births in roe deer." *Journal of Mammalogy* 74 (3): 738-744.

Gamelon, M., Besnard, A., Gaillard, J. M., Servanty, S., Baubet, E., Brandt, S. and Gimenez, O. 2011. "High hunting pressure selects for earlier birth date: wild boar as a case study." *Evolution: International Journal of Organic Evolution* 65 (11): 3100-3112.

Garnier, J. N., Holt, W. V. and Watson, P. F. 2002. "Non-invasive assessment of oestrous cycles and evaluation of reproductive seasonality in the female wild black rhinoceros (*Diceros bicornis minor*)." *Reproduction* 123 (6): 877-889.

Gogan, P. J. P., Podruzny, K. M., Olexa, E. M., Pac, H. I. and Frey, K. L. 2005. "Yellowstone bison fetal development and phenology of parturition." *The Journal of Wildlife Management* 69 (4): 1716-1730.

Gosling, L. M. 1969. "Parturition and related behaviour in Coke's hartebeest, *Alcelaphus buselaphus cokei* Günther." *Journal of Reproduction and Fertility Supplement* 6: 265-286.

Gottdenker, N. and Bodmer, R. E. 1998. "Reproduction and productivity of white-lipped and collared peccaries in the Peruvian Amazon." *Journal of Zoology* 245 (4): 423-430.

Green, W. C. H. and Rothstein, A. 1991. "Sex bias or equal opportunity? Patterns of maternal investment in bison." *Behavioral Ecology and Sociobiology* 29 (5): 373-384.

Green, W. C. H. and Rothstein, A. 1993. "Asynchronous parturition in bison: implications for the hider-follower dichotomy." *Journal of Mammalogy* 74 (4): 920-925.

Gregg, M. A., Bray, M., Kilbride, K. M. and Dunbar, M. R. 2001. "Birth synchrony and survival of pronghorn fawns." *The Journal of Wildlife Management* 65 (1): 19-24.

Grimsdell, J. J. 1973. "Reproduction in the African buffalo, *Syncerus caffer*, in western Uganda." *Journal of Reproduction and Fertility Supplement* 19: 303-318.

Guinness, F. E., Gibson, R. M. and Clutton-Brock, T. H. 1978. "Calving times of red deer (*Cervus elaphus*) on Rhum." *Journal of Zoology* 185 (1): 105-114.

Hagen, R., Ortmann, S., Elliger, A. and Arnold, J. 2021. "Advanced roe deer (*Capreolus capreolus*) parturition date in response to climate change." *Ecosphere* 12 (11): e03819.

Hart, E. E., Fennessy, J., Wells, E. and Ciuti, S. 2021. "Seasonal shifts in sociosexual behaviour and reproductive phenology in giraffe." *Behavioral Ecology and Sociobiology* 75 (1): 15.

Haskell, S. P., Ballard, W. B., Butler, D. A., Wallace, M. C., Stephenson, T. R., Alcumbrac, O. J. and Humphrey, M. H. 2008. "Factors affecting birth dates of sympatric deer in west-central Texas." *Journal of Mammalogy* 89 (2): 448-458.

Hass, C. C. 1997. "Seasonality of births in bighorn sheep." *Journal of Mammalogy* 78 (4): 1251-1260.

Hogg, J. T., Dunn, S. J., Poissant, J., Pelletier, F. and Byers, J. A. 2017. "Capital vs. income-dependent optimal birth date in two North American ungulates." *Ecosphere* 8 (4): e01766.

Hvidberg-Hansen, H. and De Vos, A. 1971. "Reproduction, population and herd structure of two Thomson's gazelle (*Gazella Thomsonii Günther*) populations." *Mammalia* 35 (1): 1-16.

Jarnemo, A., Liberg, O., Lockowandt, S., Olsson, A. and Wahlström, K. 2004. "Predation by red fox on European roe deer fawns in relation to age, sex, and birth date." *Canadian Journal of Zoology* 82 (3): 416-422.

Jongejan, G., Arcese, P. and Sinclair, A. R. E. 1991. "Growth, size and the timing of births in an individually identified population of oribi." *African Journal of Ecology* 29 (4): 340-352.

Kautz, T. M., Belant, J. L., Beyer Jr, D. E., Strickland, B. K., Petroelje, T. R. and Sollmann, R. 2019. "Predator densities and white-tailed deer fawn survival." *The Journal of Wildlife Management* 83 (5): 1261-1270.

Keech, M. A., Bowyer, R. T., Jay, M., Hoef, V., Boertje, R. D., Dale, B. W. and Stephenson, T. R. 2000. "Life-history consequences of maternal condition in Alaskan moose." *The Journal of Wildlife Management* 64 (2): 450-462.

Klingel, H. 1969. "The social organisation and population ecology of the plains zebra (*Equus quagga*)." *African Zoology* 4 (2): 249-263.

Kourkgy, C., Garel, M., Appolinaire, J., Loison, A. and Toïgo, C. 2016. "Onset of autumn shapes the timing of birth in Pyrenean chamois more than onset of spring." *Journal of Animal Ecology* 85 (2): 581-590.

Laforge, M. P., Webber, Q. M. R. and Vander Wal, E. 2023. "Plasticity and repeatability in spring migration and parturition dates with implications for annual reproductive success." *Journal of Animal Ecology* 92 (5): 1042-1054.

Lee, D. E., Bond, M. L. and Bolger, D. T. 2017. "Season of birth affects juvenile survival of giraffe." *Population Ecology* 59 (1): 45-54.

Leuthold, W. and Leuthold, B. M. 1975. "Temporal patterns of reproduction in ungulates of Tsavo East National Park." *East African Wildlife Journal* 13: 159-169.

Linnell, J. D. C. and Andersen, R. 1998. "Timing and synchrony of birth in a hider species, the roe deer *Capreolus capreolus*." *Journal of Zoology* 244 (4): 497-504.

Macchi, E., Cucuzza, A. S., Badino, P., Odore, R., Re, F., Bevilacqua, L. and Malfatti, A. 2010. "Seasonality of reproduction in wild boar (*Sus scrofa*) assessed by fecal and plasmatic steroids." *Theriogenology* 73 (9): 1230-1237.

Malmsten, A., Jansson, G., Lundeheim, N. and Dalin, A. M. 2017. "The reproductive pattern and potential of free ranging female wild boars (*Sus scrofa*) in Sweden." *Acta Veterinaria Scandinavica* 59 (1): 52.

Marshall, P. J. and Sayer, J. A. 1976. "Population ecology and response to cropping of a hippopotamus population in eastern Zambia." *Journal of Applied Ecology*: 391-403.

McGinnes, B. S. and Downing, R. L. 1977. "Factors affecting the peak of white-tailed deer fawning in Virginia." *The Journal of Wildlife Management* 41 (4): 715-719.

Meng, X., Yang, O., Feng, Z., Xia, L., Jiang, Y. and Wang, P. 2003. "Timing and synchrony of parturition in Alpine musk deer (*Moschus syfanicus*)." *Folia Zoologica* 52 (1): 39-50.

Michel, E. S., Strickland, B. K., Demarais, S., Belant, J. L., Kautz, T. M., Duquette, J. F., Beyer Jr, D. E., Chamberlain, M. J., Miller, K. V., Shuman, R. M., Kilgo, J. C., Diefenbach, D. R., Wallingford, B. D., Vreeland, J. K., Ditchkoff, S. S., DePerno, C.

S., Moorman, C. E., Chitwood, M. C. and Lashley, M. A. 2020. "Relative reproductive phenology and synchrony affect neonate survival in a nonprecocial ungulate." *Functional Ecology* 34: 2536-2547.

Moe, S. R., Rutina, L. P. and Du Toit, J. T. 2007. "Trade-off between resource seasonality and predation risk explains reproductive chronology in impala." *Journal of Zoology* 273 (3): 237-243.

Moretti, M. 1995. "Birth distribution, structure and dynamics of a hunted mountain population of wild boars (*Sus scrofa* L.), Ticino, Switzerland." *Journal of Mountain Ecology* 3: 192-196.

Moss, C. J. 2001. "The demography of an African elephant (*Loxodonta africana*) population in Amboseli, Kenya." *Journal of Zoology* 255 (2): 145-156.

Moyes, K., Nussey, D. H., Clements, M. N., Guinness, F. E., Morris, A., Morris, S., Pemberton, J. M., Kruuk, L. E. B. and Clutton-Brock, T. H. 2011. "Advancing breeding phenology in response to environmental change in a wild red deer population." *Global Change Biology* 17 (7): 2455-2469.

Ndibalema, V. G. 2009. "A comparison of sex ratio, birth periods and calf survival among Serengeti wildebeest sub-populations, Tanzania." *African Journal of Ecology* 47 (4): 574-582.

Nefdt, R. J. C. 1996. "Reproductive seasonality in Kafue lechwe antelope." *Journal of Zoology* 239 (1): 155-166.

Neumann, W., Singh, N. J., Stenbacka, F., Malmsten, J., Wallin, K., Ball, J. P. and Ericsson, G. 2020. "Divergence in parturition timing and vegetation onset in a large herbivore—differences along a latitudinal gradient." *Biology Letters* 16 (6): 20200044.

Nussey, D. H., Kruuk, L. E. B., Donald, Al, Fowlie, M. and Clutton-Brock, T. H. 2006. "The rate of senescence in maternal performance increases with early-life fecundity in red deer." *Ecology Letters* 9 (12): 1342-1350.

Ogutu, J. O., Owen-Smith, N., Piepho, H. P. and Dublin, H. T. 2015. "How rainfall variation influences reproductive patterns of African savanna ungulates in an equatorial region where photoperiod variation is absent." *PLoS One* 10 (8): e0133744.

Ogutu, J. O., Piepho, H. P. and Dublin, H. T. 2013. "Responses of phenology, synchrony and fecundity of breeding by African ungulates to interannual variation in rainfall." *Wildlife Research* 40 (8): 698-717.

Ogutu, J. O., Piepho, H. P. and Dublin, H. T. 2014. "Reproductive seasonality in African ungulates in relation to rainfall." *Wildlife Research* 41 (4): 323-342.

Ogutu, J. O., Piepho, H. P., Dublin, H. T., Bhola, N. and Reid, R. S. 2010. "Rainfall extremes explain interannual shifts in timing and synchrony of calving in topi and warthog." *Population Ecology* 52 (1): 89-102.

Ogutu, J. O., Piepho, H. P., Dublin, H. T., Bhola, N. and Reid, R. S. 2011. "Dynamics of births and juvenile recruitment in Mara–Serengeti ungulates in relation to climatic and land use changes." *Population Ecology* 53 (1): 195-213.

Owen-Smith, N. and Ogutu, J. O. 2013. "Controls over reproductive phenology among ungulates: allometry and tropical-temperate contrasts." *Ecography* 36 (3): 256-263.

Panzacchi, M., Linnell, J. D. C., Odden, J., Odden, M. and Andersen, R. 2008. "When a generalist becomes a specialist: patterns of red fox predation on roe deer fawns under contrasting conditions." *Canadian Journal of Zoology* 86 (2): 116-126.

Patterson, B. R., Mills, K. J., Middel, K. R., Benson, J. F. and Obbard, M. E. 2016. "Does predation influence the seasonal and diel timing of moose calving in central Ontario, Canada?". *PLoS One* 11 (4): e0150730.

Peláez, M., Gaillard, J. M., Bollmann, K., Heurich, M. and Rehnus, M. 2020. "Large scale variation in birth timing and synchrony of a large herbivore along the latitudinal and altitudinal gradients." *Journal of Animal Ecology* 89: 1906-1917.

Plard, F., Gaillard, J. M., Bonenfant, C., Hewison, A.J. M., Delorme, D., Cargnelutti, B., Kjellander, P., Nilsen, E. B. and Coulson, T. 2013. "Parturition date for a given female is highly repeatable within five roe deer populations." *Biology Letters* 9 (1): 20120841.

Plard, F., Gaillard, J. M., Coulson, T., Hewison, A.J. M., Delorme, D., Warnant, C. and Bonenfant, C. 2014. "Mismatch between birth date and vegetation phenology slows the demography of roe deer." *PLoS Biology* 12 (4): e1001828.

Plard, F., Gaillard, J. M., Coulson, T., Hewison, A.J. M., Delorme, D., Warnant, C., Nilsen, E. B. and Bonenfant, C. 2014. "Long-lived and heavier females give birth earlier in roe deer." *Ecography* 37 (3): 241-249.

Post, E. and Forchhammer, M. C. 2008. "Climate change reduces reproductive success of an Arctic herbivore through trophic mismatch." *Philosophical transactions of the Royal Society B: Biological sciences* 363 (1501): 2367-2373.

Post, E. and Klein, D. R. 1999. "Caribou calf production and seasonal range quality during a population decline." *The Journal of Wildlife Management* 63 (1): 335-345.

Post, E., Bøving, P. S., Pedersen, C. and MacArthur, M. A. 2003. "Synchrony between caribou calving and plant phenology in depredated and non-depredated populations." *Canadian Journal of Zoology* 81 (10): 1709-1714.

Priyadarshini, K. V. R., Gort, G., Rice, C. G. and Yoganand, K. 2022. "The reproductive phenology of blackbuck: influence of seasonal nutritional resources and flexible lactation as an adaptive strategy." *Journal of Zoology* 316 (1): 11-23.

Rachlow, J. L. and Bowyer, R. T. 1991. "Interannual Variation in Timing and Synchrony of Parturition in Dall's Sheep." *Journal of Mammalogy* 72 (3), 08, : 487-492.

Raganella-Pelliccioni, E., Scremin, M. and Toso, S. 2007. "Phenology and synchrony of roe deer breeding in northern Italy." *Acta Theriologica* 52 (1): 95-100.

Rehnus, M., Peláez, M. and Bollmann, K. 2020. "Advancing plant phenology causes an increasing trophic mismatch in an income breeder across a wide elevational range." *Ecosphere* 11 (6): e03144.

Renaud, L. A., Festa-Bianchet, M. and Pelletier, F. 2022. "Testing the match–mismatch hypothesis in bighorn sheep in the context of climate change." *Global Change Biology* 28 (1): 21-32.

Renaud, L. A., Pigeon, G., Festa-Bianchet, M. and Pelletier, F. 2019. "Phenotypic plasticity in bighorn sheep reproductive phenology: from individual to population." *Behavioral Ecology and Sociobiology* 73 (4): 50.

Rioux-Paquette, E., Festa-Bianchet, M. and Coltman, D. W. 2011. "Sex-differential effects of inbreeding on overwinter survival, birth date and mass of bighorn lambs." *Journal of Evolutionary Biology* 24 (1): 121-131.

Rodríguez-Ramírez, M. and Mora, J. M. 2022. "Analysis of the male annual antler cycle, reproductive behavior and spotted fawn presence in the tropical white-tailed deer." *Therya* 13 (2): 143-151.

Rosser, A. M. 1989. "Environmental and reproductive seasonality of puku, *Kobus vardoni*, in Luangwa Valley, Zambia." *African Journal of Ecology* 27 (1): 77-88.

Rubin, E. S., Boyce, W. M. and Bleich, V. C. 2000. "Reproductive strategies of desert bighorn sheep." *Journal of Mammalogy* 81 (3): 769-786.

Rutberg, A. T. 1984. "Birth synchrony in American bison (*Bison bison*): response to predation or season?". *Journal of Mammalogy* 65 (3): 418-423.

Rutberg, A. T. 1987. "Adaptive hypotheses of birth synchrony in ruminants: an interspecific test." *The American Naturalist* 130 (5): 692-710.

Ryan, S. J., Knechtel, C. U. and Getz, W. M. 2007. "Ecological cues, gestation length, and birth timing in African buffalo (*Syncerus caffer*)." *Behavioral Ecology* 18 (4): 635-644.

San José, C. and Braza, F. 1992. "Antipredator aspects of fallow deer behaviour during calving season at Donana National Park (Spain)." *Ethology Ecology & Evolution* 4 (2): 139-149.

Santos, P., Fernández-Llario, P., Fonseca, C., Monzón, A., Bento, P., Soares, A. M. V. M., Mateos-Quesada, P. and Petrucci-Fonseca, F. 2006. "Habitat and reproductive phenology of wild boar (*Sus scrofa*) in the western Iberian Peninsula." *European Journal of Wildlife Research* 52 (3): 207-212.

Sigouin, D., Ouellet, J. P. and Courtois, R. 1997. "Geographical variation in the mating and calving periods of moose." *Alces* 33: 85-95.

Sinclair, A. R. E., Mduma, S. A. R. and Arcese, P. 2000. "What determines phenology and synchrony of ungulate breeding in Serengeti?" *Ecology* 81 (8): 2100-2111.

Smuts, G. L. 1976. "Reproduction in the zebra mare *Equus burchelli antiquorum* from the Kruger National Park." *Koedoe* 19 (1): 89-132.

Spinage, C. A. 1969. "Reproduction in the Uganda defassa waterbuck, *Kobus defassa ugandae* Neumann." *Journal of Reproduction and Fertility* 18 (3): 445-457.

Srivastava, T., Kumar, A., Kumar, V. and Umapathy, G. 2021. "Diet drives differences in reproductive synchrony in two sympatric mountain ungulates in the Himalaya." *Frontiers in Ecology and Evolution* 9: 647465.

Stoner, D. C., Sexton, J. O., Nagol, J., Bernales, H. H. and Edwards Jr, T. C. 2016. "Ungulate reproductive parameters track satellite observations of plant phenology across latitude and climatological regimes." *PLoS One* 11 (2): e0148780.

Stopher, K. V., Pemberton, J. M., Clutton-Brock, T. H. and Coulson, T. 2008. "Individual differences, density dependence and offspring birth traits in a population of red deer." *Proceedings of the Royal Society B: Biological Sciences* 275 (1647): 2137-2145.

Testa, J. W. 2002. "Does predation on neonates inherently select for earlier births?". *Journal of Mammalogy* 83 (3): 699-706.

Testa, J. W., Becker, E. F. and Lee, G. R. 2000. "Temporal patterns in the survival of twin and single moose (*Alces alces*) calves in southcentral Alaska." *Journal of Mammalogy* 81 (1): 162-168.Thompson, R. W. and Turner, J. C. 1982. "Temporal geographic variation in the lambing season of bighorn sheep." *Canadian Journal of Zoology* 60 (8): 1781-1793.

Walling, C. A., Nussey, D. H., Morris, A., Clutton-Brock, T. H., Kruuk, L. E. B. and Pemberton, J. M. 2011. "Inbreeding depression in red deer calves." *BMC Evolutionary Biology* 11 (1): 1-13.

Walther, F. R. 1972. "Social grouping in Grant's gazelle (*Gazella granti* Brooke 1827) in the Serengeti National Park." *Zeitschrift für Tierpsychologie* 31 (4): 348-403.

Wittemyer, G., Barner Rasmussen, H. and Douglas-Hamilton, I. 2007. "Breeding phenology in relation to NDVI variability in free-ranging African elephant." *Ecography* 30 (1): 42-50.

**Supplementary Material 3: Quality index according to time, continent and taxonomic family**

The score for each variable tested corresponds to the sum of the scores (between 1 and 3) for four categories: data collection method, sample size, study period length and type of test. The score of the article corresponds to the mean of the scores per variable tested. The higher the score (between 4 and 12), the higher the quality or the article.

Regarding data collection methods, we broadly identified 10 methods in the retrieved articles. First, we considered camera trapping (i.e. age determination of juveniles based on camera trap images), faecal sample (i.e. dosage of reproductive hormones in female's faeces to estimate parturition date) and aerial survey (i.e. identification of the age and number of juveniles from an aircraft) to be the less reliable methods. Inaccurate assessment of the age of the juvenile from a distance or from a limited number of pictures with important risks of multiple observations of the same individual can lead to biased distributions of births. Similarly, assessing the time of parturition from hormonal dosage of faecal samples is bound to show huge uncertainty. We thus attributed the score of 1 to these methods.

Second, we considered that extrapolation of the date of birth from foetus size or from the weight of culled individuals using growth curves was more reliable than the previous methods, but still affected by inter-annual and individual variability (under-nutrition can slow down foetus growth; e.g., Roine et al. 1982; Holmes et al. 2021). Similarly, age identification of juveniles based on direct observations in the field, which can vary from daily observations of individually-known specimens to transect counts among large groups of unidentified individuals, was also entailed by a large uncertainty. These methods were consequently attributed the intermediary score of 2. The "literature review" method corresponded to datasets extracted by the authors of a review article from several original articles. As the methods used to determine the phenology of births can vary from one original article to the other and vary in terms of reliability and precision, we also attributed the intermediary score of 2 to the method “literature review”, to account for such a variability in the articles concerned.

Finally, tracking female movement (i.e. close, often daily, monitoring to identify clusters of points at the location of parturition, usually combined with a field visit to confirm the birth), vaginal implant transmitter (i.e. electronic device placed in the vagina of the female and signalling the timing of parturition when expelled from the reproductive tract) and capture of neonate (i.e. usually no more than a few days after birth, and allowing age determination at hand) were given the highest score of 3. Although these methods present their own biases, they were considered as the most reliable to estimate the date of birth of juveniles in our study, obtained either directly at hand or by using precise temporal analysis of female parturition time.

Table S3.1: Categories used to design the article quality index and the associated score per level for each category (data collection method, sample size, study period length, type of test).

| **Category** | **Level** | **Score** |
|---|---|---|
| **Data collection method** | Camera trapping | 1 |
| | Faecal sample | 1 |
| | Aerial survey | 1 |
| | Culled specimen | 2 |
| | Foetus | 2 |
| | Direct observations | 2 |
| | Literature review | 2 |
| | Movement tracking | 3 |
| | Vaginal implant transmitter | 3 |
| | Capture of neonate | 3 |
| **Sample size** | ]0; 25] | 1 |
| | ]25; 100] | 2 |
| | >100 | 3 |
| **Study period length** | 1 year | 1 |
| | 2-5 years | 2 |
| | >5 years | 3 |
| **Type of test** | Descriptive | 1 |
| | Statistical | 3 |

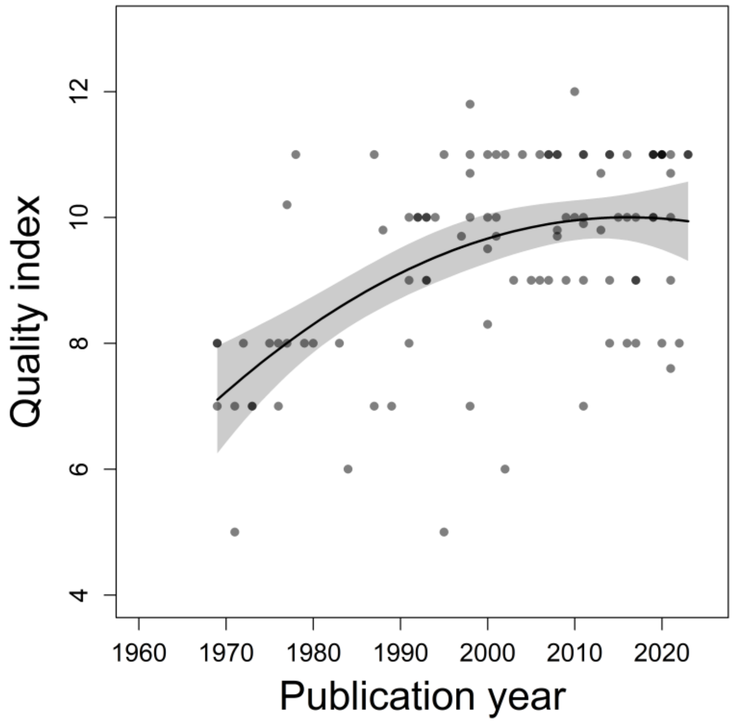


Figure S3.1: Change in the quality index of the articles investigating the drivers of the phenology of birth in large herbivores between 1960 and 2024 (n = 103 articles with a calculable index). The dots represent the quality index of each article, the line represents the predictions of the quadratic regression, the shaded area represents the 95% confidence interval.

Table S3.1: Output of the post-hoc test (Tukey Honest Significant Differences) investigating the effect of continent on quality index of the articles (n = 103 articles).

| **Groups compared** | **Diffe-rence** | **Lower interval** | **Upper interval** | **p** |
|---|---|---|---|---|
| Asia *vs.* Africa | -0.45 | -2.35 | 1.44 | 0.96 |
| Central_and_South_America *vs.* Africa | -0.35 | -2.25 | 1.54 | 0.99 |
| Europe *vs.* Africa | 1.98 | 1.04 | 2.92 | 0.00 |
| North_America *vs.* Africa | 1.60 | 0.73 | 2.46 | 0.00 |
| Central_and_South_America *vs.* Asia | 0.10 | -2.42 | 2.62 | 1.00 |
| Europe *vs.* Asia | 2.44 | 0.53 | 4.35 | 0.01 |
| North_America *vs.* Asia | 2.05 | 0.17 | 3.93 | 0.02 |
| Europe *vs.* Central_and_South_America | 2.34 | 0.43 | 4.25 | 0.01 |
| North_America *vs.* Central_and_South_America | 1.95 | 0.07 | 3.83 | 0.04 |
| North_America *vs.* Europe | -0.39 | -1.29 | 0.52 | 0.76 |

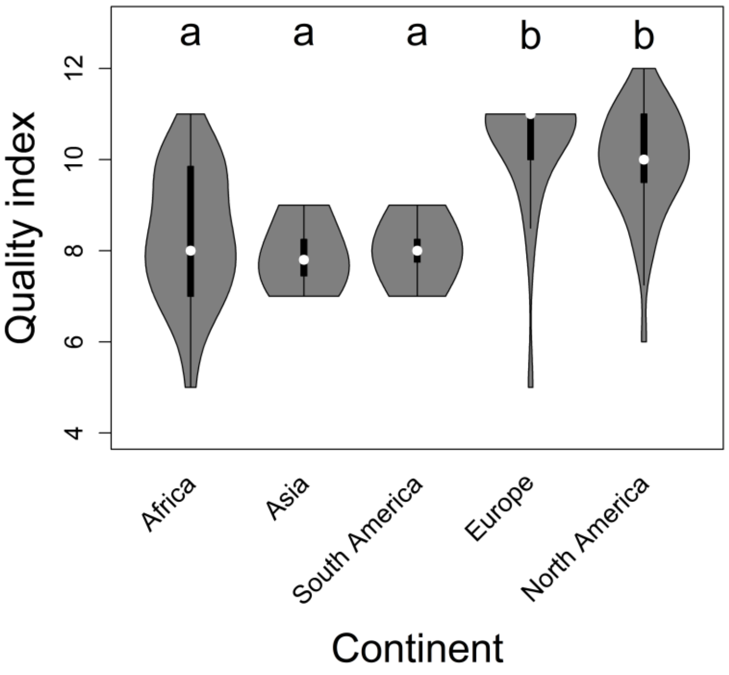


Figure S3.2: Quality index of the articles investigating the drivers of the phenology of birth in large herbivores between 1960 and 2024 (n = 103 articles with a calculable index) according to the continent of origin of the populations studied (in the figure, "South America" stands for "Central and South America"). The letters represent significant differences between groups.

Table S3.2: Output of the post-hoc test (Tukey Honest Significant Differences) investigating the effect of taxonomic family on quality index of the articles (n = 103 articles).

| **Groups compared** | **Diffe-rence** | **Lower interval** | **Upper interval** | **p** |
|---|---|---|---|---|
| Cervidae-Moschidae *vs.* Suidae-Tayassuidae | 1.75 | 0.61 | 2.89 | 0.00 |
| Equidae *vs.* Suidae-Tayassuidae | 0.04 | -1.67 | 1.75 | 1.00 |
| Bovidae-Antilocapridae *vs.* Suidae-Tayassuidae | 0.19 | -0.92 | 1.30 | 1.00 |
| Giraffidae *vs.* Suidae-Tayassuidae | 0.34 | -1.37 | 2.05 | 0.99 |
| Elephantidae-Hippopotamidae-Rhinocerotidae *vs.* Suidae-Tayassuidae | -0.15 | -2.06 | 1.77 | 1.00 |
| Equidae *vs.* Cervidae-Moschidae | -1.71 | -3.19 | -0.24 | 0.01 |
| Bovidae-Antilocapridae *vs.* Cervidae-Moschidae | -1.57 | -2.26 | -0.87 | 0.00 |
| Giraffidae *vs.* Cervidae-Moschidae | -1.41 | -2.89 | 0.06 | 0.07 |
| Elephantidae-Hippopotamidae-Rhinocerotidae *vs.* Cervidae-Moschidae | -1.90 | -3.61 | -0.19 | 0.02 |
| Bovidae-Antilocapridae *vs.* Equidae | 0.15 | -1.30 | 1.60 | 1.00 |
| Giraffidae *vs.* Equidae | 0.30 | -1.65 | 2.25 | 1.00 |
| Elephantidae-Hippopotamidae-Rhinocerotidae *vs.* Equidae | -0.19 | -2.32 | 1.95 | 1.00 |
| Giraffidae *vs.* Bovidae-Antilocapridae | 0.15 | -1.30 | 1.60 | 1.00 |
| Elephantidae-Hippopotamidae-Rhinocerotidae *vs.* Bovidae-Antilocapridae | -0.33 | -2.02 | 1.36 | 0.99 |
| Elephantidae-Hippopotamidae-Rhinocerotidae *vs.* Giraffidae | -0.49 | -2.62 | 1.65 | 0.99 |

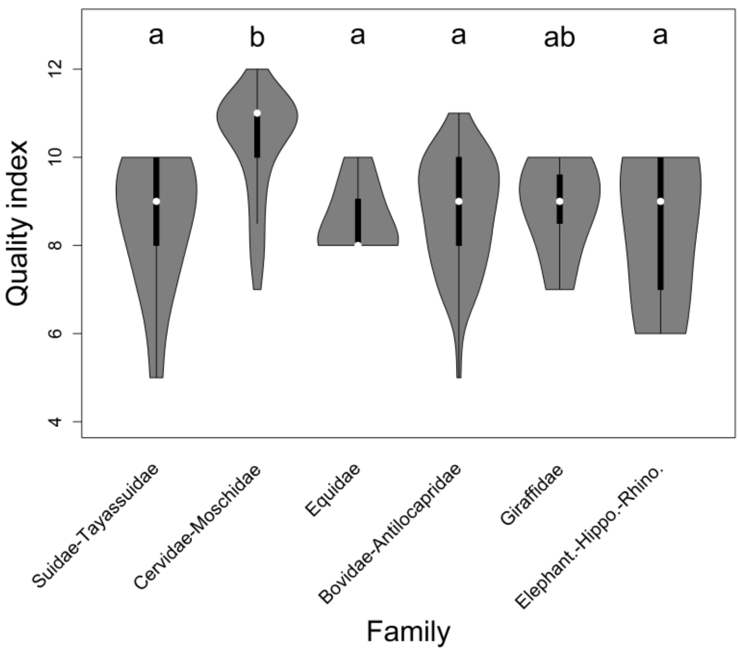


Figure S3.3: Quality index of the articles investigating the drivers of the phenology of birth in large herbivores between 1960 and 2024 (n = 103 articles with a calculable index) according to the taxonomic family of the populations studied (in the figure, "Elephant.-Hippo.-Rhino." stands for "Elephantidae-Hippopotamidae-Rhinocerotidae"). The letters represent significant differences between groups.

**Supplementary Material 4: Distribution of factors per theme**

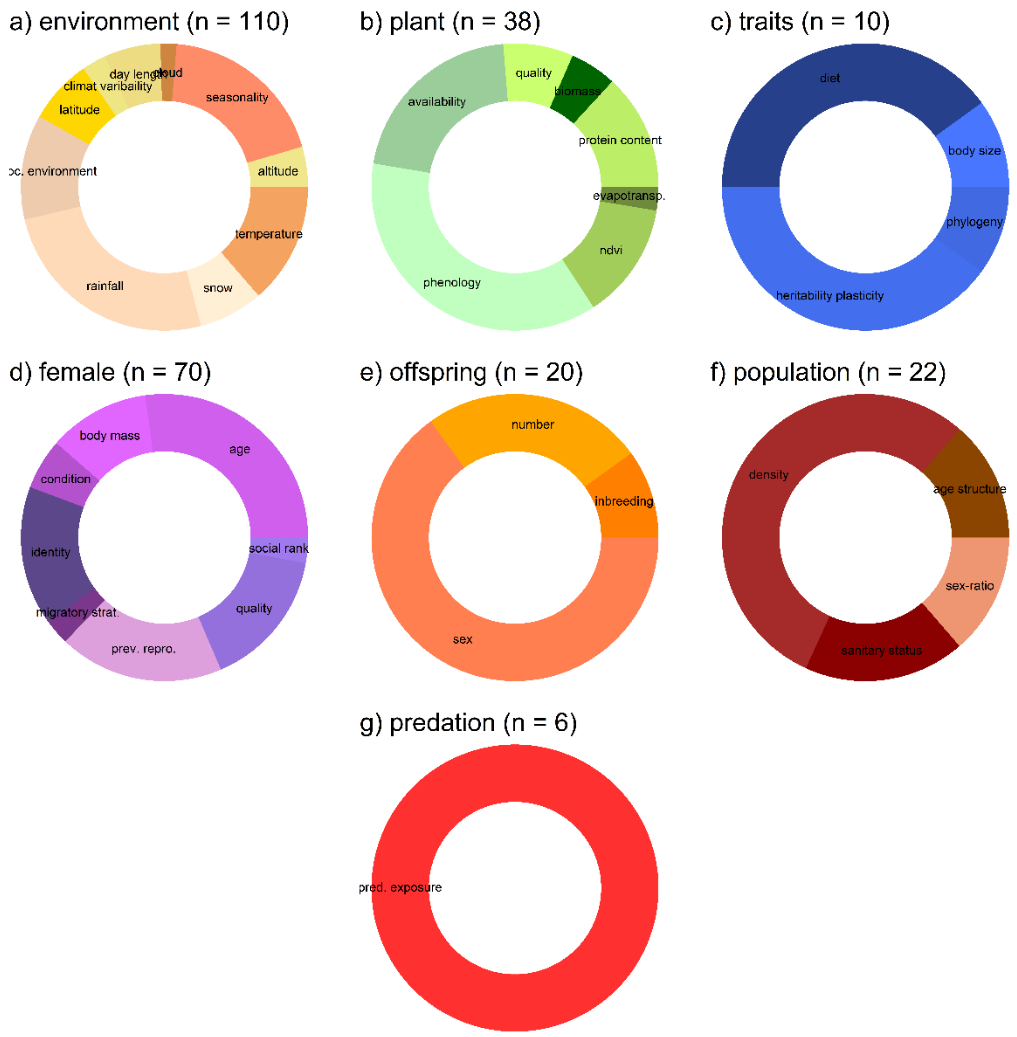


Figure S4.1: Distribution of the factors investigated for their effect on the timing of birth in large herbivores for each theme (n = 7 themes; environmental characteristics; plant characteristics; life history traits; female characteristics; offspring characteristics; population characteristics; predation exposure) in the articles retained in the study (published between 1960 and 2024). Each figure corresponds to the number of articles investigating the effect of a given factor at least once. Abbreviations: loc. environment stands for local environmental conditions, evapostransp. stands for plant evapotranspiration, prev. repro. stands for previous reproductive status, migratory strat. stands for migratory strategies, pred. exposure stands for predation exposure, anti-pred. behav. stands for anti-predator behaviour of the juvenile.

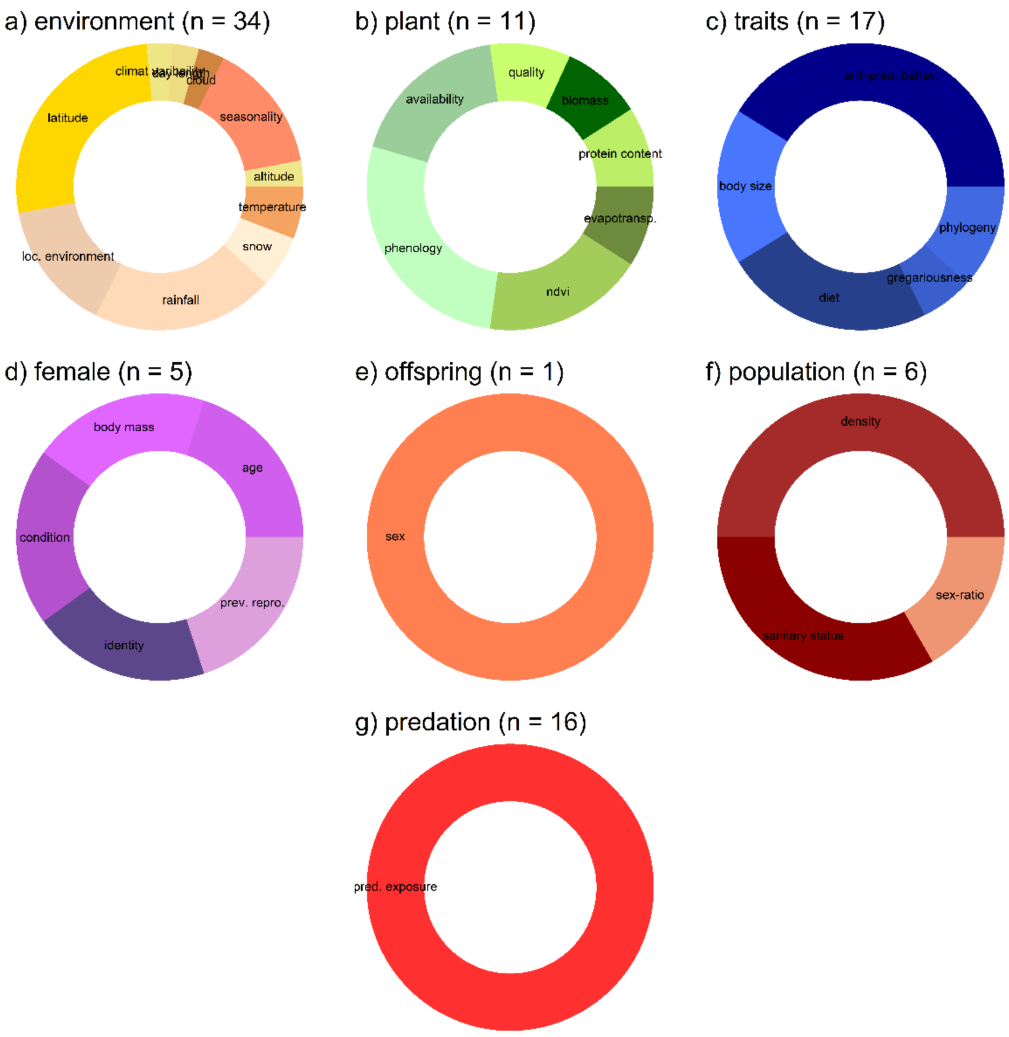


Figure S4.2: Distribution of the factors investigated for their effect on the synchrony of birth in large herbivores for each theme (n = 7 themes; environmental characteristics; plant characteristics; life history traits; female characteristics; offspring characteristics; population characteristics; predation exposure) in the articles retained in the study (published between 1960 and 2024). Each figure corresponds to the number of articles investigating the effect of a given factor at least once. Abbreviations: loc. environment stands for local environmental conditions, evapostransp. stands for plant evapotranspiration, prev. repro. stands for previous reproductive status, migratory strat. stands for migratory strategies, pred. exposure stands for predation exposure, anti-pred. behav. stands for anti-predator behaviour of the juvenile.

**Supplementary Material 5: Number of times each factor tested received support, partial support and no support**

Table S5.1: Number of times a factor is tested for its effect on the timing of births in large herbivores, and receives no support, partial support or support.

| Theme | Factor | No support | Partial support | Support | Total |
|---|---|---|---|---|---|
| environmental characteristics | altitude | 1 | 0 | 6 | 7 |
| environmental characteristics | annual climatic seasonality | 7 | 13 | 27 | 47 |
| environmental characteristics | cloud | 2 | 0 | 0 | 2 |
| environmental characteristics | day length | 2 | 1 | 6 | 9 |
| environmental characteristics | inter-annual climatic variability | 2 | 0 | 1 | 3 |
| environmental characteristics | latitude | 4 | 0 | 8 | 12 |
| environmental characteristics | local environmental conditions | 2 | 2 | 9 | 13 |
| environmental characteristics | rainfall | 19 | 3 | 41 | 63 |
| environmental characteristics | snow | 5 | 0 | 4 | 9 |
| environmental characteristics | temperature | 8 | 1 | 14 | 23 |
| plant characteristics | fecal or forage protein content | 11 | 0 | 8 | 19 |
| plant characteristics | forage biomass | 11 | 0 | 2 | 13 |
| plant characteristics | forage quality | 2 | 0 | 1 | 3 |
| plant characteristics | general forage availability | 3 | 0 | 5 | 8 |
| plant characteristics | general plant phenology | 4 | 2 | 13 | 19 |
| plant characteristics | NDVI | 2 | 0 | 4 | 6 |
| plant characteristics | plant evapotranspiration | 0 | 0 | 2 | 2 |
| life history traits | body size | 0 | 1 | 0 | 1 |
| life history traits | diet | 6 | 1 | 4 | 11 |
| life history traits | heritability - plasticity | 1 | 0 | 3 | 4 |
| life history traits | phylogeny | 0 | 0 | 1 | 1 |
| female characteristics | female age | 9 | 1 | 10 | 20 |
| female characteristics | female body mass | 2 | 1 | 7 | 10 |

| Theme | Factor | No support | Partial support | Support | Total |
|---|---|---|---|---|---|
| female characteristics | female condition | 2 | 1 | 2 | 5 |
| female characteristics | female identity | 3 | 2 | 10 | 15 |
| female characteristics | female migratory strategies | 1 | 0 | 1 | 2 |
| female characteristics | female previous reproductive status | 5 | 1 | 9 | 15 |
| female characteristics | female quality | 5 | 3 | 6 | 14 |
| female characteristics | female social rank | 2 | 0 | 0 | 2 |
| offspring characteristics | offspring inbreeding | 1 | 0 | 1 | 2 |
| offspring characteristics | offspring number | 3 | 0 | 2 | 5 |
| offspring characteristics | offspring sex | 11 | 1 | 2 | 14 |
| population characteristics | population age structure | 2 | 0 | 1 | 3 |
| population characteristics | population density | 6 | 0 | 6 | 12 |
| population characteristics | population sanitary state | 0 | 2 | 2 | 4 |
| population characteristics | population sex ratio | 2 | 0 | 1 | 3 |
| predation exposure | predation exposure | 2 | 1 | 4 | 7 |

Table S5.2: Number of times a factor is tested for its effect on the synchrony of births in large herbivores, and receives no support, partial support or support.

| **Theme** | **Factor** | **No support** | **Partial support** | **Support** | **Total** |
|---|---|---|---|---|---|
| environmental characteristics | altitude | 0 | 0 | 1 | 1 |
| environmental characteristics | annual climatic seasonality | 1 | 1 | 9 | 11 |
| environmental characteristics | cloud | 1 | 0 | 0 | 1 |
| environmental characteristics | day length | 0 | 0 | 1 | 1 |
| environmental characteristics | inter-annual climatic variability | 0 | 0 | 1 | 1 |
| environmental characteristics | latitude | 5 | 0 | 5 | 10 |
| environmental characteristics | local environmental conditions | 3 | 1 | 3 | 7 |
| environmental characteristics | rainfall | 13 | 8 | 3 | 24 |
| environmental characteristics | snow | 3 | 0 | 0 | 3 |
| environmental characteristics | temperature | 6 | 0 | 0 | 6 |
| plant characteristics | fecal or forage protein content | 5 | 0 | 0 | 5 |
| plant characteristics | forage biomass | 8 | 4 | 1 | 13 |
| plant characteristics | forage quality | 1 | 0 | 0 | 1 |
| plant characteristics | general forage availability | 0 | 0 | 2 | 2 |
| plant characteristics | general plant phenology | 2 | 1 | 1 | 4 |
| plant characteristics | NDVI | 0 | 0 | 2 | 2 |
| plant characteristics | plant evapotranspiration | 0 | 0 | 1 | 1 |
| life history traits | anti-predator behaviour of the juvenile | 7 | 0 | 11 | 18 |
| life history traits | body size | 1 | 0 | 2 | 3 |
| life history traits | diet | 2 | 1 | 1 | 4 |
| life history traits | gregariousness | 1 | 0 | 0 | 1 |
| life history traits | phylogeny | 0 | 1 | 1 | 2 |
| female characteristics | female age | 0 | 0 | 2 | 2 |
| female characteristics | female body mass | 1 | 0 | 0 | 1 |
| female characteristics | female condition | 0 | 0 | 1 | 1 |
| female characteristics | female identity | 1 | 0 | 1 | 2 |

| Theme | Factor | No support | Partial support | Support | Total |
|---|---|---|---|---|---|
| female characteristics | female previous reproductive status | 0 | 0 | 2 | 2 |
| offspring characteristics | offspring sex | 0 | 0 | 1 | 1 |
| population characteristics | population density | 5 | 0 | 4 | 9 |
| population characteristics | population sanitary state | 2 | 0 | 0 | 2 |
| population characteristics | population sex ratio | 1 | 0 | 0 | 1 |
| predation exposure | predation exposure | 12 | 1 | 7 | 20 |

**Supplementary Material 6: Number of times an article, a species, or a population is used to assess the strength of the effect of a given theme on timing and synchrony**

Table S6.1: Number of times an article is used to assess the effect of a given theme on timing and synchrony. Only articles appearing more than once are reported.

| Phenology parameter | Theme | Nb repetitions | Article reference |
|---|---|---|---|
| timing | environmental characteristics | 2 - 5 | Bowyer 1998; Bunnell 1982; Dasmann 1962; Froy 2019; Gottdenker 1998; Grimsdell 1973; Hart 2021; Haskell 2008; Hass 1997; Hvidberg-Hansen 1971; Klingel 1969; Kourkgy 2016; Lee 2017; Leuthold 1975; McGinnes 1977; Meng 2003; Moe 2007; Moyes 2011; Neumann 2020; Ogutu 2010; Ogutu 2015; Owen-Smith 2013; Pelaez 2020; Plard 2014b; Renaud 2019; Renaud 2022; Smuts 1976; Srivastava 2021; Stoner 2016; Thompson 1982 |
| timing | environmental characteristics | 6 - 10 | Fairall 1968; Ogutu 2011; Ogutu 2014; Sigouin 1997 |
| timing | environmental characteristics | > 10 | Ogutu 2013; Sinclair 2000 |
| timing | plant characteristics | 2 - 5 | Aikens 2021; Bunnell 1982; Chen 2018; Feder 2008; Field 1973; Hogg 2017; Post 2007; Priyadarshini 2022; Renaud 2022; Thompson 1982 |
| timing | plant characteristics | > 10 | Sinclair 2000 |
| timing | female characteristics | 2 - 5 | Adams 1998; Aikens 2021; Berger 1992; Bonnet 2019; Cote 2001; Feder 2008; Festa-Bianchet 1988; Flydal 2002; Green 1991; Guinness 1978; Haskell 2008; Kourkgy 2016; Laforge 2023; Linnell 1998; Moyes 2011; Nussey 2006; Renaud 2019; Rubin 2000; Stopher 2008; Walling 2011 |
| timing | female characteristics | 6 - 10 | Plard 2013; Plard 2014a |
| timing | offspring characteristics | 2 - 5 | Festa-Bianchet 1988; Linnell 1998; Rioux-Paquette 2011 |
| timing | pop. characteristics | 2 - 5 | Bonnet 2019; Festa-Bianchet 1988; Kourkgy 2016 |
| timing | life history traits | 2 - 5 | Owen-Smith 2013; Sinclair 2000; Srivastava 2021 |
| timing | life history traits | 6 - 10 | Ogutu 2014 |
| timing | predation exposure | 2 - 5 | Panzacchi 2008 |
| synchrony | environmental characteristics | 2 - 5 | Bowyer 1998; Couriot 2023; English 2012; Linnell 1998; Moe 2007; Ogutu 2010; Ogutu 2014; Pelaez 2020 |
| synchrony | environmental characteristics | 6 - 10 | Ogutu 2013; Sigouin 1997 |
| synchrony | environmental characteristics | > 10 | Sinclair 2000 |
| synchrony | plant characteristics | 2 - 5 | Thompson 1982 |
| synchrony | plant characteristics | > 10 | Sinclair 2000 |
| synchrony | female characteristics | 2 - 5 | Berger 1992; Hogg 2017; Neumann 2020 |
| synchrony | pop. characteristics | 6 - 10 | Sinclair 2000 |
| synchrony | life history traits | 2 - 5 | English 2012; Ogutu 2013; Ogutu 2014; Rutberg 1987 |
| synchrony | life history traits | > 10 | Sinclair 2000 |
| synchrony | predation exposure | 2 - 5 | Estes 1976; Michel 2020 |

Table S6.2: Number of times a species is used to assess the effect of a given theme on timing and synchrony. Only species appearing more than once are reported.

| **Phenology parameter** | **Theme** | **Nb repetitions** | **Species** |
|---|---|---|---|
| timing | environmental characteristics | 2 - 5 | *Antilocapra americana; Bison bison; Bison bonasus; Bos mutus; Camelus ferus; Capra ibex; Ceratotherium simum; Cervus albirostis; Cervus canadensis; Cervus nippon; Connochaetes gnou; Connochaetes taurinus; Dama dama; Damaliscus pygargus; Diceros bicornis; Equus ferus; Equus kiang; Eudorcas thomsonii; Hemitragus jemlahicus; Hippopotamus amphibius; Hippotragus equinus; Hippotragus niger; Kobus kob; Kobus leche; Kobus vardonii; Loxodonta africana; Madoqua kirkii; Moschus chrysogaster; Nanger granti; Oreamnos americanus; Oryx gazella; Ourebia ourebi; Ovibos moschatus; Ovis ammon; Ovis gmelini; Ovis musimon; Ovis vignei; Pantholops hodgsonii; Pecari tajacu; Pelea capreolus; Phacochoerus aethiopicus; Procapra gutturosa; Procapra picticaudata; Pseudois nayaur; Redunca fulvorufula; Rupicapra pyrenaica; Rupicapra rupicapra; Saiga tatarica; Sus scrofa; Taurotragus oryx; Tragelaphus angasii; Tragelaphus buxtoni; Tragelaphus scriptus; Tragelaphus strepsiceros* |
| timing | environmental characteristics | 6 - 10 | *Alcelaphus buselaphus; Cervus elaphus; Kobus ellipsiprymnus; Odocoileus hemionus; Ovis dalli; Phacochoerus africanus; Rangifer tarandus; Syncerus caffer* |
| timing | environmental characteristics | > 10 | *Aepyceros melampus; Alces alces; Capreolus capreolus; Damaliscus lunatus; Equus quagga; Giraffa camelopardalis; Odocoileus virginianus; Ovis canadensis* |
| timing | plant characteristics | 2 - 5 | *Aepyceros melampus; Alcelaphus buselaphus; Alces alces; Antilope cervicapra; Capreolus capreolus; Connochaetes taurinus; Damaliscus lunatus; Equus quagga; Eudorcas thomsonii; Giraffa camelopardalis; Kobus ellipsiprymnus; Nanger granti; Odocoileus hemionus; Ourebia ourebi; Ovis dalli; Phacochoerus aethiopicus; Syncerus caffer* |
| timing | plant characteristics | 6 - 10 | *Rangifer tarandus* |
| timing | plant characteristics | > 10 | *Ovis canadensis* |
| timing | female characteristics | 2 - 5 | *Alces alces; Bison bison; Odocoileus hemionus; Odocoileus virginianus; Rupicapra pyrenaica* |
| timing | female characteristics | 6 - 10 | *Rangifer tarandus* |
| timing | female characteristics | > 10 | *Capreolus capreolus; Cervus elaphus; Ovis canadensis* |
| timing | offspring characteristics | 2 - 5 | *Alces alces; Bison bison; Capreolus capreolus; Cervus elaphus; Odocoileus hemionus* |
| timing | offspring characteristics | 6 - 10 | *Ovis canadensis* |
| timing | population characteristics | 2 - 5 | *Rupicapra pyrenaica* |
| timing | population characteristics | 6 - 10 | *Cervus elaphus; Ovis canadensis* |
| timing | life history traits | 2 - 5 | *Aepyceros melampus; Alcelaphus buselaphus; Alces alces; Antilocapra americana; Bison bison; Bison bonasus; Bos mutus; Camelus ferus; Capra ibex; Capreolus capreolus; Ceratotherium simum; Cervus albirostis; Cervus canadensis; Cervus elaphus; Cervus nippon; Connochaetes gnou; Connochaetes taurinus;* |

| Phenology parameter | Theme | Nb repetitions | Species |
|---|---|---|---|
| | | | *Dama dama; Damaliscus lunatus; Damaliscus pygargus; Diceros bicornis; Equus ferus; Equus kiang; Equus quagga; Eudorcas thomsonii; Giraffa camelopardalis; Hemitragus jemlahicus; Hippopotamus amphibius; Hippotragus equinus; Hippotragus niger; Kobus ellipsiprymnus; Kobus kob; Kobus leche; Kobus vardonii; Loxodonta africana; Madoqua kirkii; Nanger granti; Odocoileus hemionus; Odocoileus virginianus; Oreamnos americanus; Oryx gazella; Ourebia ourebi; Ovibos moschatus; Ovis ammon; Ovis canadensis; Ovis dalli; Ovis gmelini; Ovis musimon; Ovis vignei; Pantholops hodgsonii; Pelea capreolus; Phacochoerus aethiopicus; Phacochoerus africanus; Procapra gutturosa; Procapra picticaudata; Pseudois nayaur; Rangifer tarandus; Redunca fulvorufula; Rupicapra rupicapra; Saiga tatarica; Syncerus caffer; Taurotragus oryx; Tragelaphus angasii; Tragelaphus buxtoni; Tragelaphus scriptus; Tragelaphus strepsiceros* |
| timing | predation exposure | 2 - 5 | *Alces alces; Capreolus capreolus* |
| synchrony | environmental characteristics | 2 - 5 | *Alcelaphus buselaphus; Antilocapra americana; Bison bison; Cervus canadensis; Cervus elaphus; Connochaetes taurinus; Damaliscus pygargus; Equus quagga; Eudorcas thomsonii; Giraffa camelopardalis; Hippotragus niger; Kobus ellipsiprymnus; Kobus kob; Kobus leche; Madoqua kirkii; multiple; Nanger granti; Odocoileus hemionus; Odocoileus virginianus; Oreamnos americanus; Ourebia ourebi; Ovis canadensis; Ovis dalli; Phacochoerus aethiopicus; Phacochoerus africanus; Rangifer tarandus; Redunca fulvorufula; Syncerus caffer; Tragelaphus angasii; Tragelaphus scriptus; Tragelaphus strepsiceros* |
| synchrony | environmental characteristics | 6 - 10 | *Aepyceros melampus; Capreolus capreolus; Damaliscus lunatus* |
| synchrony | environmental characteristics | > 10 | *Alces alces* |
| synchrony | plant characteristics | 2 - 5 | *Aepyceros melampus; Alcelaphus buselaphus; Alces alces; Bison bison; Connochaetes taurinus; Damaliscus lunatus; Eudorcas thomsonii; Giraffa camelopardalis; Kobus ellipsiprymnus; Nanger granti; Odocoileus hemionus; Ourebia ourebi; Ovis canadensis; Rangifer tarandus; Syncerus caffer* |
| synchrony | female characteristics | 2 - 5 | *Alces alces; Antilocapra americana; Bison bison; Ovis canadensis* |
| synchrony | population characteristics | 2 - 5 | *Bison bison; Connochaetes taurinus* |
| synchrony | life history traits | 2 - 5 | *Alces alces; Antilocapra americana; Bison bison; Cervus canadensis; Cervus elaphus; Connochaetes taurinus; Damaliscus pygargus; Equus quagga; Eudorcas thomsonii; Hippotragus niger; Kobus kob; Kobus leche; multiple; Nanger granti; Odocoileus hemionus; Odocoileus virginianus; Oreamnos americanus; Ovis aries; Ovis dalli; Phacochoerus africanus; Philantomba monticola; Rangifer tarandus; Redunca fulvorufula; Syncerus caffer; Tragelaphus angasii; Tragelaphus scriptus; Tragelaphus strepsiceros* |
| synchrony | life history traits | 6 - 10 | *Aepyceros melampus; Alcelaphus buselaphus; Damaliscus lunatus; Giraffa camelopardalis* |
| synchrony | predation exposure | 2 - 5 | *Alces alces; Capreolus capreolus; Connochaetes taurinus; Odocoileus virginianus; Rangifer tarandus* |

Table S6.3: Number of times a population is used to assess the effect of a given theme on timing and synchrony. Only populations appearing more than once are reported.

| **Phenology parameter** | **Theme** | **Nb repetitions** | **Populations** |
|---|---|---|---|
| timing | environmental characteristics | 2 - 5 | *Rangifer tarandus*-Alaska; *Capreolus capreolus*-Storfosna; *Cervus elaphus*-Isle of Rum; *Alces alces*-Denali National Park; *Capreolus capreolus*-Trois Fontaines; *Giraffa camelopardalis*-Northern Namib Desert; *Eudorcas thomsonii*-Serengeti National Park; *Ourebia ourebi*-Serengeti National Park; *Equus quagga*-Serengeti National Park; *Rupicapra pyrenaica*-Bazes; *Giraffa camelopardalis*-Tarangire Ecosystem; *Capreolus capreolus*-Europe; *Odocoileus virginianus*-Radford Army Ammunition Plant; *Moschus chrysogaster*-Xinglongshan National Nature Reserve; *Aepyceros melampus*-Chobe National Park; *Phacochoerus aethiopicus*-Masai Mara National Reserve; *Alcelaphus buselaphus*-Masai Mara National Reserve; *Equus quagga*-Masai Mara National Reserve; *Aepyceros melampus*-Masai Mara National Reserve; *Giraffa camelopardalis*-Masai Mara National Reserve; *Phacochoerus africanus*-Masai Mara National Reserve; *Capreolus capreolus*-Alps; *Nanger granti*-Serengeti National Park; *Odocoileus hemionus*-White Mountains to Wasatch Mountains; *Ovis dalli*-North America; *Odocoileus hemionus*-West Central Texas; *Odocoileus virginianus*-West Central Texas |
| timing | environmental characteristics | 6 - 10 | *Equus quagga*-Kruger National Park; *Ovis canadensis*-Ram Mountain; *Damaliscus lunatus*-Masai Mara National Reserve; *Ovis canadensis*-North America; *Alces alces*-Ontario; *Alces alces*-Newfoundland; *Alces alces*-Wyoming; *Alces alces*-Alaska; *Alces alces*-British Columbia; *Alces alces*-Yukon; *Alces alces*-Quebec; *Alces alces*-Montana |
| timing | environmental characteristics | > 10 | *Alces alces*-Sweden |
| timing | plant characteristics | 2 - 5 | *Odocoileus hemionus*-Rocky Mountains; *Rangifer tarandus*-Bathurst; *Ovis canadensis*-Ram Mountain; *Eudorcas thomsonii*-Serengeti National Park; *Ourebia ourebi*-Serengeti National Park; *Equus quagga*-Serengeti National Park; *Connochaetes taurinus*-Serengeti National Park; *Rangifer tarandus*-Kangerlussuaq; *Antilope cervicapra*-Velavadar National Park; *Nanger granti*-Serengeti National Park; *Damaliscus lunatus*-Serengeti National Park; *Syncerus caffer*-Serengeti National Park; *Giraffa camelopardalis*-Serengeti National Park; *Aepyceros melampus*-Serengeti National Park; *Kobus ellipsiprymnus*-Serengeti National Park; *Phacochoerus aethiopicus*-Serengeti National Park; *Alcelaphus buselaphus*-Serengeti National Park; *Ovis canadensis*-North America; *Ovis dalli*-North America |
| timing | female characteristics | 2 - 5 | *Rangifer tarandus*-Denali National Park; *Odocoileus hemionus*-Rocky Mountains; *Bison bison*-Badlands National Park; *Ovis canadensis*-Caw Ridge; *Ovis canadensis*-Sheep River Drainage; *Cervus elaphus*-Ilse of Rum; *Bison bison*-Wind Cave National Park; *Rupicapra pyrenaica*-Bazes; *Rangifer tarandus*-Newfoundland; *Rangifer tarandus*-North Ottadalen; *Rangifer tarandus*-South Ottadalen; *Rangifer tarandus*-Snohetta; *Ovis canadensis*-Bradley Canyon; *Ovis canadensis*-Deep Canyon; *Ovis canadensis*-San Ysidro Mountains; *Ovis canadensis*-Carrizo Canyon; *Odocoileus hemionus*-West Central Texas; *Odocoileus virginianus*-West Central Texas |

| Phenology parameter | Theme | Nb repetitions | Populations |
|---|---|---|---|
| timing | female characteristics | 6 - 10 | *Capreolus capreolus*-Storfosna; *Ovis canadensis*-Ram Mountain; *Capreolus capreolus*-Aurignac; *Capreolus capreolus*-Bogesund; *Capreolus capreolus*-Grimso |
| timing | female characteristics | > 10 | *Cervus elaphus*-Isle of Rum; *Capreolus capreolus*-Trois Fontaines |
| timing | offspring characteristics | 2 - 5 | *Capreolus capreolus*-Storfosna; *Cervus elaphus*-Isle of Rum; *Ovis canadensis*-Ram Mountain; *Ovis canadensis*-Sheep River Drainage |
| timing | population characteristics | 2 - 5 | *Ovis canadensis*-Ram Mountain; *Ovis canadensis*-Sheep River Drainage; *Rupicapra pyrenaica*-Bazes |
| timing | population characteristics | 6 - 10 | *Cervus elaphus*-Isle of Rum |
| timing | life history traits | 2 - 5 | *Eudorcas thomsonii*-Serengeti National Park; *Ourebia ourebi*-Serengeti National Park; *Equus quagga*-Serengeti National Park; *Connochaetes taurinus*-Serengeti National Park; *Nanger granti*-Serengeti National Park; *Damaliscus lunatus*-Serengeti National Park; *Syncerus caffer*-Serengeti National Park; *Giraffa camelopardalis*-Serengeti National Park; *Aepyceros melampus*-Serengeti National Park; *Kobus ellipsiprymnus*-Serengeti National Park; *Phacochoerus aethiopicus*-Serengeti National Park; *Madoqua kirkii*-Serengeti National Park; *Alcelaphus buselaphus*-Serengeti National Park |
| synchrony | environmental characteristics | 2 - 5 | *Rangifer tarandus*-Alaska; *Alces alces*-Denali National Park; *Eudorcas thomsonii*-Serengeti National Park; *Capreolus capreolus*-Europe; *Aepyceros melampus*-Chobe National Park; *Aepyceros melampus*-Mala Mala Private Game Reserve; *Connochaetes taurinus*-Serengeti National Park; *Phacochoerus aethiopicus*-Masai Mara National Reserve; *Damaliscus lunatus*-Masai Mara National Reserve; *Alcelaphus buselaphus*-Masai Mara National Reserve; *Aepyceros melampus*-Masai Mara National Reserve; *Phacochoerus africanus*-Masai Mara National Reserve; *Bison bison*-National Bison Range; *Nanger granti*-Serengeti National Park; *Damaliscus lunatus*-Serengeti National Park; *Syncerus caffer*-Serengeti National Park; *Giraffa camelopardalis*-Serengeti National Park; *Aepyceros melampus*-Serengeti National Park; *Alcelaphus buselaphus*-Serengeti National Park; *Rangifer tarandus*-Northern Canada; *Rangifer tarandus*-Western Alaska |
| synchrony | environmental characteristics | 6 - 10 | *Alces alces*-Sweden; *Alces alces*-Ontario; *Alces alces*-Newfoundland; *Alces alces*-Wyoming; *Alces alces*-British Columbia; *Alces alces*-Yukon; *Alces alces*-Quebec; *Alces alces*-Montana |
| synchrony | environmental characteristics | > 10 | *Alces alces*-Alaska |
| synchrony | plant characteristics | 2 - 5 | *Eudorcas thomsonii*-Serengeti National Park; *Ourebia ourebi*-Serengeti National Park; *Connochaetes taurinus*-Serengeti National Park; *Nanger granti*-Serengeti National Park; *Damaliscus lunatus*-Serengeti National Park; *Syncerus caffer*-Serengeti National Park; *Giraffa camelopardalis*-Serengeti National Park; *Aepyceros melampus*-Serengeti National Park; *Kobus ellipsiprymnus*-Serengeti National Park; *Alcelaphus buselaphus*-Serengeti National Park; *Ovis canadensis*-North America |
| synchrony | female characteristics | 2 - 5 | *Bison bison*-Badlands National Park; *Ovis canadensis*-National Bison Range; *Antilocapra americana*-National Bison Range; *Alces alces*-Sweden |
| synchrony | population characteristics | 2 - 5 | *Connochaetes taurinus*-Serengeti National Park |

| Phenology parameter | Theme | Nb repetitions | Populations |
|---|---|---|---|
| synchrony | life history traits | 2 - 5 | *Rangifer tarandus*-Alaska; *Tragelaphus scriptus*-Central Rift Province; *Tragelaphus angasii*-Ndumu Reserve; *Cervus elaphus*-Isle of Rum; *Connochaetes taurinus*-Ngorongoro Crater; *Antilocapra americana*-National Bison Range; *Eudorcas thomsonii*-Serengeti National Park; *Connochaetes taurinus*-Serengeti National Park; *Damaliscus lunatus*-Masai Mara National Reserve; *Alcelaphus buselaphus*-Masai Mara National Reserve; *Equus quagga*-Masai Mara National Reserve; *Aepyceros melampus*-Masai Mara National Reserve; *Giraffa camelopardalis*-Masai Mara National Reserve; *Phacochoerus africanus*-Masai Mara National Reserve; Bison bison-National Bison Range; *Nanger granti*-Serengeti National Park; *Damaliscus lunatus*-Serengeti National Park; *Syncerus caffer*-Serengeti National Park; *Giraffa camelopardalis*-Serengeti National Park; *Aepyceros melampus*-Serengeti National Park; *Alcelaphus buselaphus*-Serengeti National Park; *Ovis dalli*-Sheep Mt; *Ovis aries*-Saint Kilda; *Oreamnos americanus*-Mount Wardle; *Cervus canadensis*-North Central Michigan; *Alces alces*-Trondelag; *Odocoileus virginianus*-South Texas; *Odocoileus hemionus*-Western Oregon; *Rangifer tarandus*-Northwestern Alaska; *Kobus kob*-Rwenzori National Park; *Kobus leche*-Kafue Flats; *Redunca fulvorufula*-Loskop Dam; *Redunca fulvorufula*-Gilgil; *Alcelaphus buselaphus*-Central Rift Province; *Damaliscus pygargus*-Bontebok National Park; *Damaliscus pygargus*-Van Riebeeck Reserve; *Damaliscus lunatus*-Moremi Reserve; *Aepyceros melampus*-Sengwa Reserve; *Hippotragus niger*-Shimba Hills Reserve; *Tragelaphus strepsiceros*-Guluene Chefu Reserve; *Nanger granti*-Central Rift Province; *Eudorcas thomsonii*-Central Rift Province; *Philantomba monticola*-Tsitsikamma Coastal National Park; *Giraffa camelopardalis*-Tibavati Reserve; *Alces alces*-Alaska |
| synchrony | predation exposure | 2 - 5 | *Connochaetes taurinus*-Ngorongoro Crater; *Odocoileus virginianus*-Fort Rucker; *Odocoileus virginianus*-Tensas River National Wildlife Refuge; *Odocoileus virginianus*-Escanaba; *Odocoileus virginianus*-Crystal Falls; *Odocoileus virginianus*-Fort Bragg; *Odocoileus virginianus*-Brosnan Forrest; *Odocoileus virginianus*-Savannah River; *Odocoileus virginianus*-Penns Valley; *Odocoileus virginianus*-Quehanna Wild Area |
